# Dose-dependent ultrasound neuromodulation of the human brain

**DOI:** 10.64898/2026.09.27.754841

**Authors:** Benjamin R. Kop, Morteza Mohammadjavadi, Eva Feredoes, Martin T.W. Scott, Anthony M. Norcia, Elsa Fouragnan, Ryan T. Ash, Gary H. Glover, Kim Butts Pauly

## Abstract

Transcranial ultrasound stimulation (TUS) can noninvasively modulate deep brain structures, but the relationship between delivered dose and neuromodulatory response remains poorly characterized in humans. Here, we show that ultrasound-induced tissue displacement measured in vivo predicts local and downstream functional neuromodulation. In 24 healthy adults, we combined magnetic resonance acoustic radiation force imaging (MR-ARFI) with concurrent TUS-fMRI in a double-blind, counterbalanced, within-subject design. Relative to active control stimulation, targeting the lateral geniculate nucleus (LGN) increased visually evoked BOLD responses in the LGN and ipsilateral primary visual cortex, localized to the retinotopic projection zone of the stimulated LGN site. Displacement predicted effect magnitude in both regions across participants and covaried with voxelwise responses within participants. Simulated in situ intensity showed no significant association with displacement or neuromodulatory response. These findings link in vivo mechanical displacement to functional modulation of a human thalamocortical circuit, providing an empirical basis for individualized dosing in causal neuroscience and clinical intervention.

## Introduction

Transcranial ultrasound stimulation (TUS) promises to transform causal neuroscientific research and clinical intervention through precision neuromodulation anywhere in the brain^1^. In little over a decade, human studies have progressed from early demonstrations^2^ to causal neuroscientific insights^3–5^ and encouraging clinical effects^6^. However, the field has not yet delineated the relationship between the dose of ultrasound delivered and neuromodulation effects^7,8^. This is likely a major contributor to the inter-individual and inter-study variability in neuromodulatory outcomes, which together form a major obstacle to realising the full potential of TUS.

Heterogeneity in TUS dose arises from two major sources: idiosyncrasies in the skull and variability in acoustic coupling^9^, both of which are difficult to constrain at the individual participant level. Biophysical modeling of transcranial ultrasound propagation currently carries considerable uncertainty in amplitude estimates^10^, and coupling quality is left unmodelled. As a result, simulations are limited in their ability to resolve dose-response relationships, leaving the link between transcranial dose and neuromodulatory effects poorly characterized.

Magnetic resonance acoustic radiation force imaging (MR-ARFI) offers a solution^11^. This imaging technique phase encodes the tissue displacement elicited by TUS acoustic radiation force, providing an absorption- and tissue-mechanics-weighted metric of in vivo dose. While MR-ARFI has been applied in non-human primates^12,13^, and recently as a standalone method in humans^14^, inter-individual variability in displacement and its influence on functional outcomes has remained unquantified. Here, we demonstrate the utility of MR-ARFI^14^ for targeting and in vivo dosimetry in human neuromodulation research, using a concurrent TUS-fMRI protocol.

We build on prior work targeting the lateral geniculate nucleus of the thalamus (LGN)^11,15–17^, which provides a clear paradigm to evaluate dose-dependent circuit engagement. This core visual node is small, deep, lateralized, and contralaterally organized. It projects monosynaptically to the downstream visual cortex through a well-characterized retinotopic pathway. This architecture permits both local and network-level readouts of dose-dependent functional engagement, thus providing a readily controlled model for fundamental neuroscientific investigations of brain-wide circuitry, as well as a sensory analog of the pathological circuit neuromodulation desired for neuropsychiatric intervention^18,19^. Based on prior human and animal work, we hypothesized that TUS would increase visual-system BOLD responses^16^ and that the magnitude of this increase would scale with empirically quantified in vivo displacement^11,20^.

## Results

We delivered TUS to unilateral LGN and measured effects on task-related blood oxygen level-dependent (BOLD) fMRI in 24 healthy participants (M_age_ = 28.3 ± 7.5 years, range 19-49; 14 male) using a within-subjects, double-blind, counterbalanced, online TUS-fMRI design (Fig. 1A). A 64-element phased array was used for steered 3D targeting of the LGN in-scanner (Fig. 1B). The LGN was localized structurally and functionally using a fMRI localizer processed in real time. Contrast-reversing polar checkerboards were used to evoke visual activation in LGN and visual cortices. Group-level analysis confirmed that the visual stimuli elicited robust lateralized activation in both the LGN and visual cortex (Fig. 1C). This task-related activation was then modulated with LGN-TUS. MR-ARFI scans were acquired to quantify in-vivo displacement (Fig. 1D).

**Fig. 1:**
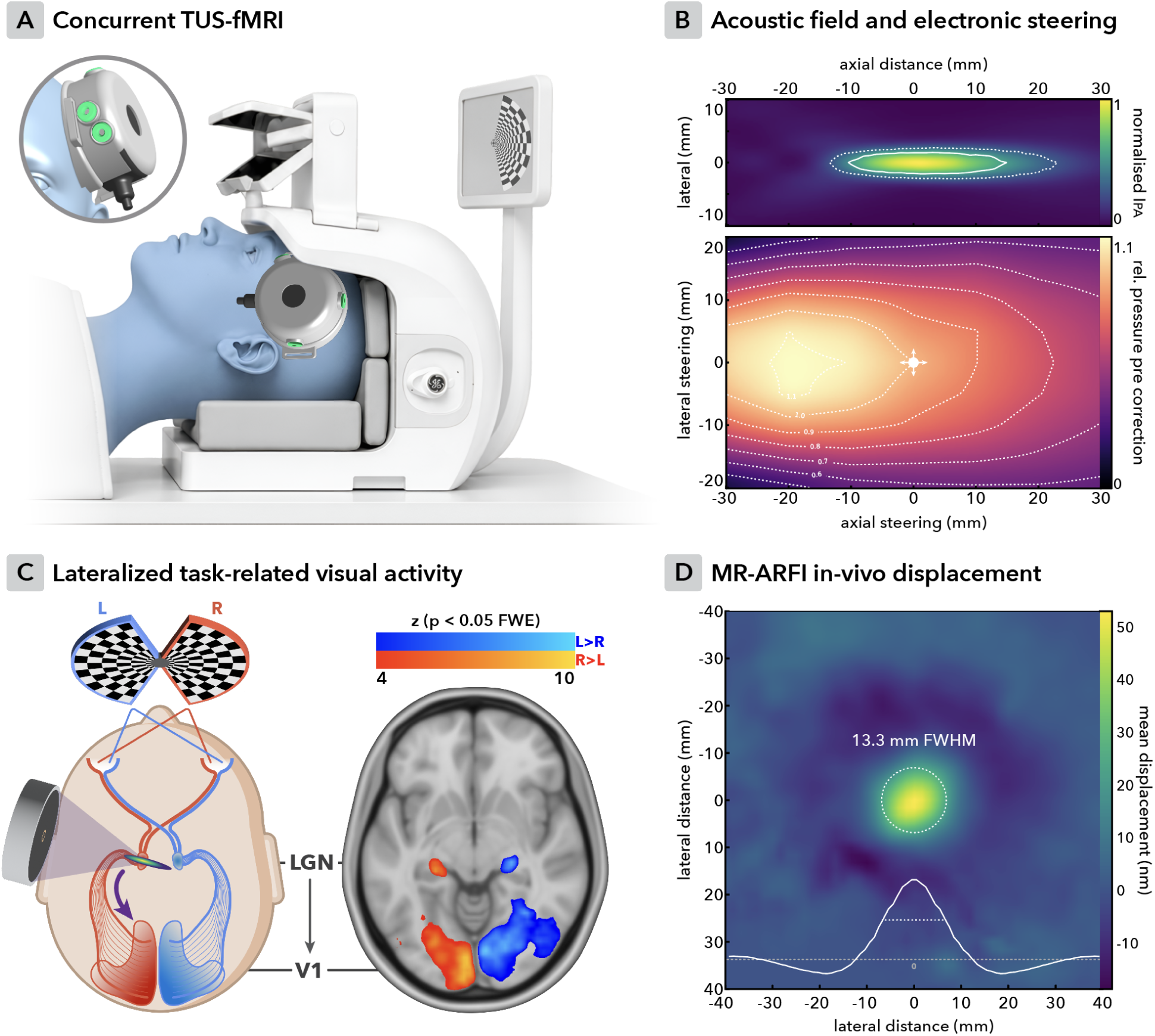
Experimental design. **(A)** Concurrent TUS-fMRI design using an MR-compatible 64-channel phased array transducer placed anterosuperior to the left ear and directed at the LGN with a 10° posterior-inferior angled custom holder. Fiducials (dark grey) were used for transducer reconstruction and phased acoustic steering. **(B)** Top: acoustic field at the geometric focus without electronic steering in normalized pulse-average intensity (I_PA_). The geometric focus was at a 75 mm axial depth with an axial full-width-half-maximum intensity of 25 mm (FWHM; solid line = -3 dB; dotted line = -6 dB). Bottom: relative pressure as compared to pressure without electronic steering across the lateral and axial range. These data were used to correct the drive amplitude such that identical free-water TUS pressure/intensity was delivered for all steering settings. **(C)** We studied dose-dependent effects of TUS on BOLD fMRI visual activation evoked by hemifield contrast-reversing polar checkerboards. Left: Schematic of the experiment showing participants viewing hemifield contrast-reversing polar checkerboards. The visual system is lateralized such that right hemifield visual drive (red) passes through left LGN to left V1. Sonication of the left LGN is therefore expected to specifically increase task-related activation locally in the left LGN and on the network-level in downstream V1, without affecting left hemifield / right hemisphere responses (blue). Right: empirical data from the main experiment (N = 24) depicting lateralized visual activation in both the LGN and primary visual cortices. **(D)** The average MR-ARFI displacement map across subjects reveals a well-defined focus with an in-plane FWHM of 13.3 mm (N = 21).

We tested an active control site TUS (temporal grey matter) and a baseline condition with no sonication to isolate direct neuromodulatory effects (Fig. 2A). Explicit assessment of blinding confirmed that participants could not reliably distinguish between LGN- and Control-TUS (Fig. 2A; t(23) = −0.11, p = 0.916; MΔ = −0.521 [95% CI: −10.7, 9.62]; d_z_ = 0.02; BF_01_ = 4.63). Sonication was administered at 500 kHz, 50 W/cm^2^ free-water I_SPPA_ for 145 ms as a continuous pulse that repeated six times per 15-second block (Fig. 2B). We hypothesized that LGN-TUS would increase this activation, in line with prior work^16^, where stronger in vivo displacement quantified with MR-ARFI would lead to stronger effects^7,11,20^. Indeed, LGN-TUS produced a spatially specific increase in local task-related neural activation that propagated along the geniculostriate pathway to the ipsilateral visual cortex. The magnitude of this increase scaled with in vivo displacement (Fig. 2C-F).

**Fig. 2:**
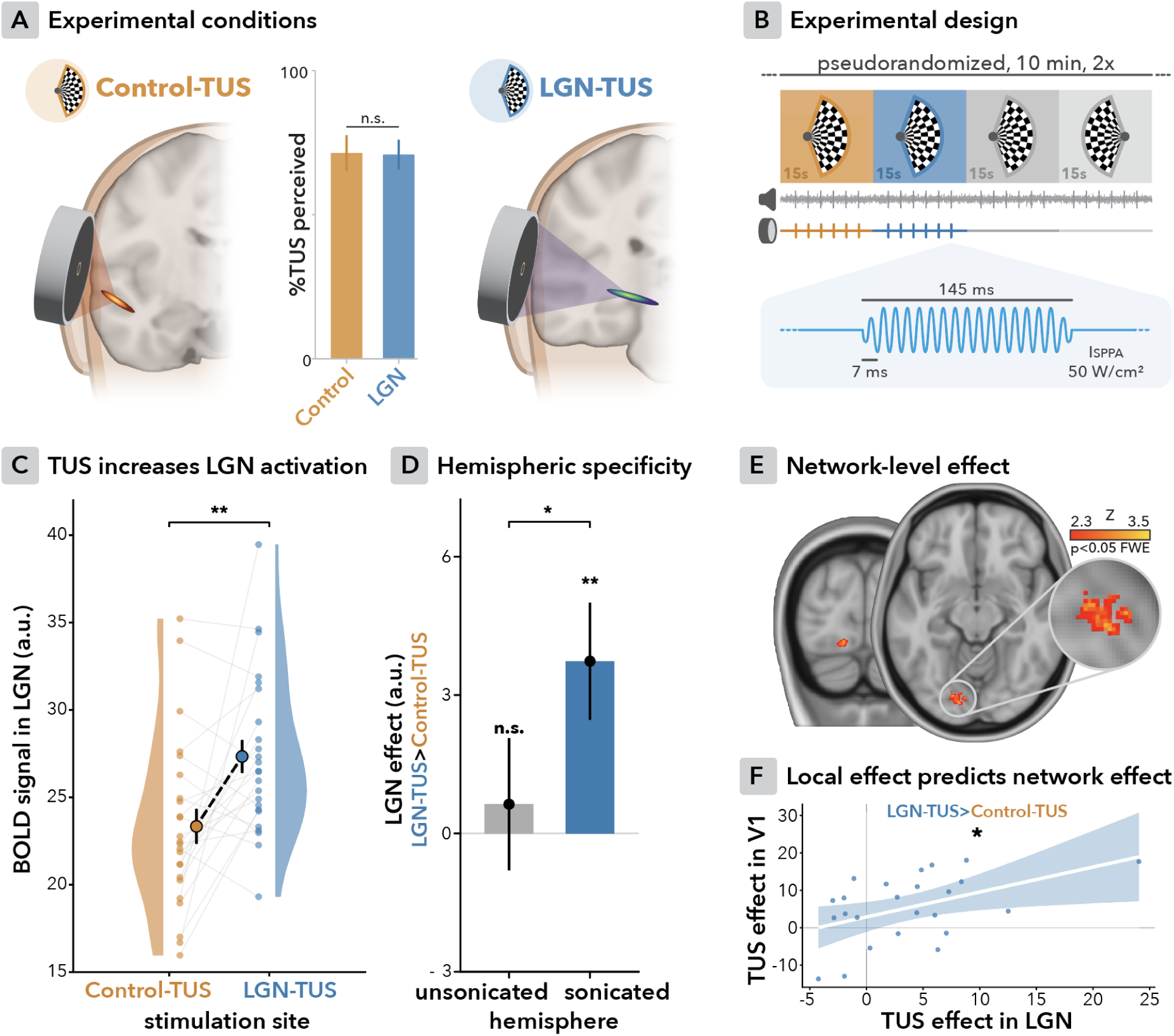
LGN-TUS increases local and network-level task-related activation. **(A)** To isolate direct neuromodulatory effects of LGN-TUS from nonspecific peripheral confounds, Control-TUS was delivered to the gray matter of the temporal lobe. This region is anatomically distinct and unrelated to visual processing. Blinding between these two primary conditions was successful (N = 24). **(B)** Conditions were tested in two 10-minute acquisitions using a counterbalanced, blocked, double-blind design. Per 15-second block, LGN-/Control-TUS was delivered six times. White noise was played through earbuds continuously during scanning. A 4 kHz tone was added during TUS (or when TUS would have been delivered during baseline conditions). Baseline conditions (grey) allowed for cooling and visualization of lateralized activation. Sonication was delivered as 145 ms, 500 kHz continuous pulses with a 7 ms ramp at 50 W/cm^2^ in an online design. **(C)** LGN-TUS significantly increases local BOLD signal compared to Control-TUS. **(D)** LGN-TUS-specific increase in BOLD signal was restricted to the sonicated LGN (left), and was absent in the unsonicated contralateral LGN. **(E)** Group-level analysis similarly showed increased BOLD signal in ipsilateral V1, restricted to the stimulated hemisphere. Cluster at z > 2.3, p (FWE) < 0.05. **(F)** The magnitude of TUS effects in the V1 cluster was significantly predicted by the magnitude of TUS effects locally in the LGN. All empirical plots depict N = 24.

### TUS increases local and network-level task-related activation

LGN-TUS increased task-related activation in the stimulated LGN relative to the active control site (Control-TUS) (Fig. 2C; t(23) = 3.12, p = 0.005, MΔ = 4.00 [95% CI: 1.34, 6.65], d_z_ = 0.64). The LGN-TUS>Control-TUS effect was significantly larger in the stimulated left LGN as compared to the unsonicated contralateral LGN (Fig. 2D; t(23) = 2.13, p = 0.044, MΔ = 3.10 [95% CI: 0.084, 6.12], d_z_ = 0.43). Correspondingly, there was no significant LGN-TUS effect in the contralateral, unsonicated LGN (Fig. 2D; t(23) = 0.44, p = 0.662, mean = 0.633 [95% CI: -2.32, 3.59], d = 0.09, BF01 = 4.26). Together, these results demonstrate spatially specific, online ultrasound engagement of a deep-brain nucleus that had previously been inferred based on downstream effects.

We additionally observed increased BOLD signal at the network level in monosynaptically-connected ipsilateral visual cortex. Group-level analysis revealed a circumscribed cluster of significantly increased task-related activation during LGN-TUS relative to Control-TUS (Fig. 2E, cluster-corrected p = 0.015), corroborating prior work^16^. No such change occurred in the contralateral visual cortex, further supporting the spatial specificity of LGN-TUS effects.

Local and network-level effects scaled together (Fig. 2F). Specifically, the magnitude of the LGN-TUS>Control-TUS effect in V1 was significantly predicted by the TUS effect size locally in the LGN (Fig. 2F; β = 0.47, 95% CI [0.08, 0.86], t(22) = 2.47, p = 0.022, R² = 0.22). Taken together, these results are in line with a spatially specific direct thalamic effect propagating transsynaptically along the geniculostriate pathway.

### MR-ARFI-quantified dose predicts functional effects

We hypothesized that MR-ARFI quantified dose would predict the size of functional response to TUS, as it has in animal work^11,20^. Indeed, greater focal in vivo displacement predicted a larger LGN BOLD signal increase during LGN-TUS as compared to control (Fig. 3A; β = 0.54, 95% CI [0.13, 0.94], t(19) = 2.77, p = 0.006 (one-sided), R² = 0.29). This same displacement likewise predicted the effect in downstream ipsilateral V1 (Fig. 3A; β = 0.47, 95% CI [0.04, 0.89], t(19) = 2.30, p = 0.016 (one-sided), R² = 0.22). These relationships also held when taking the maximum displacement in LGN specifically as the predictor (LGN: β = 0.43, 95% CI [-0.01, 0.86], t(19) = 2.05, p = 0.027 (one-sided), R² = 0.18; V1: β = 0.38, 95% CI [-0.07, 0.82], t(19) = 1.77, p = 0.046 (one-sided), R² = 0.14).

**Fig. 3:**
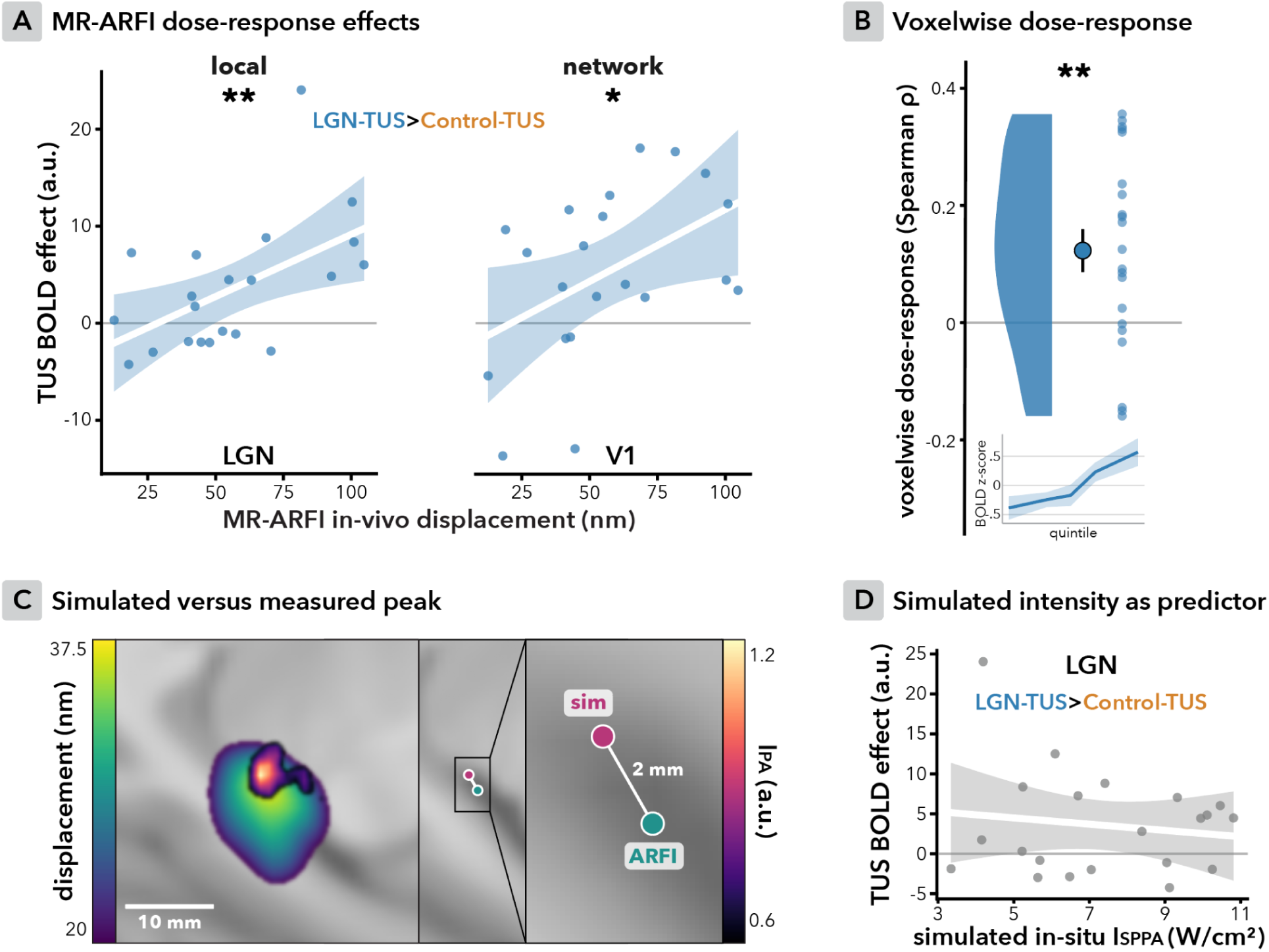
MR-ARFI-quantified dose-response effects. **(A)** As hypothesized, greater MR-ARFI-quantified in vivo displacement predicted a larger TUS effect (i.e., LGN-TUS>Control-TUS BOLD contrast), both locally in LGN (left) and downstream in ipsilateral V1 (right). **(B)** Within participants, voxels receiving greater displacement showed a larger LGN-TUS>Control-TUS effect. This is revealed by our sample of subject-level Spearman correlation coefficients being significantly greater than zero. The inset depicts normalized LGN-TUS effect by within-participant displacement quintiles, illustrating the shape of the relationship. **(C)** Simulated pulse-average intensity (I_PA_) and measured displacement peaks correspond on group-level, with only a 2 mm deviation, falling within the error range of our fiducial registration approach. **(D)** Simulated spatial-peak pulse-average intensity (I_SPPA_) did not significantly predict LGN-TUS effects (nor displacement itself), underscoring the utility of MR-ARFI in explaining meaningful inter-individual variability in the functional outcome of ultrasound neuromodulation. N = 21.

A parsimonious account of the dose-response observed in V1 is that its activation was driven by increased LGN activation via the monosynaptic geniculostriate projection. A mediation analysis tested this (Supplementary Fig. 2; displacement → LGN effect → V1 effect; 5,000 bootstrap resamples). Displacement predicted the LGN effect (path a: β = 0.43, 95% CI [0.04, 0.90]), which in turn predicted the V1 response independently of displacement (path b: β = 0.43, 95% CI [0.01, 0.96]). The direct displacement → V1 path was not significant with LGN in the model (c′: β = 0.19, 95% CI [−0.15, 0.68]). The indirect effect through LGN, i.e., the formal test of mediation, was positive, albeit with the 95% CI marginally including zero (a × b: β = 0.18, 95% CI [−0.02, 0.47]). Given our sample size, the bootstrap intervals are wide. Moreover, measurement error in a mediator (here, LGN BOLD signal) is known to attenuate the indirect path and inflate the direct path, reallocating variance from a × b to c′ without altering the total effect^21^. Therefore, we treat this analysis as consistent with, but not formal evidence of, geniculostriate dose-dependent propagation.

We additionally explored whether dose-dependence held within participants, across voxels. In the LGN itself, the voxelwise dose-response was not significant (t(20) = −0.31, p = 0.756; MΔ = −0.035 [95% CI: −0.269, 0.199]; d_z_ = −0.07). This is unsurprising when considering that the LGN is small, and therefore spans a restricted displacement range and contributes few voxels at the individual participant level, leaving little internal gradient to detect and limited sensitivity with which to detect it. Therefore, we repeated the analysis in a larger region, namely the empirically derived average full-width half-maximum (FWHM) area of displacement (i.e., the 13.3 mm average diameter of in-plane half-max; see Fig. 1D). This provided roughly 5x as many voxels per participant across a wider range of displacement. Here, voxels receiving greater displacement indeed showed a larger LGN-TUS > Control-TUS effect (Fig. 3B; t(20) = 3.41, p = 0.003; mean = 0.128 [95% CI: 0.050, 0.207]; d_z_ = 0.74).

Taken together, MR-ARFI displacement predicted the magnitude of the TUS effect both across participants and within participants. These results represent the first demonstration in humans that an in vivo, MR-ARFI-quantified metric of actualized dose can predict the functional magnitude of ultrasound neuromodulation.

#### MR-ARFI localizes the acoustic focus in vivo

To estimate dose in vivo, we acquired MR-ARFI displacement images in an oblique plane through the LGN, perpendicular to the axis of acoustic propagation. The group-average displacement map revealed a well-defined focus with a lateral in-plane FWHM of 13.3 mm (Fig. 1D; see Supplementary Fig. 1 for individual plots). The median offset between each participant’s displacement center of gravity and the LGN centroid was 5.32 mm (IQR: 3.42-7.66 mm). Despite this bias, the median LGN displacement was 98% of each participant’s peak (IQR: 62.6-100%), and significant displacement was induced directly in the LGN for 90% of participants (peak voxelwise t > 2.3).

#### Acoustic simulations localize the focus but do not predict in vivo dose or response

Unlike empirically verified displacement, simulated in-situ intensity did not significantly explain inter-individual variability in neuromodulatory response (Fig. 3D; β = −0.14, 95% CI [−0.61, 0.34], t(19) = −0.61, p = 0.550, R² = 0.02), nor did it predict the magnitude of displacement itself (Supplementary Fig. 3; β = 0.13, 95% CI [−0.35, 0.61], t(19) = 0.57, p = 0.288 (one-sided), R² = 0.02).

Importantly, simulations did estimate the location of peak displacement well. The discrepancy between group-average MR-ARFI and simulated peaks was only ∼2 mm (Fig. 3C), roughly on the same order as our transducer fiducial registration error (Supplementary Fig. 4). These results corroborate prior reports that simulations carry meaningful uncertainty in transcranial estimates, here demonstrated in vivo.

### Local and network-level effects are retinotopically concordant

To further explore the connection between local and network-level effects, we investigated whether the most strongly exposed region of the LGN corresponded retinotopically to the significant cluster of V1 activation. The group-level standard space (MNI152) average displacement map revealed targeting concentrated in the posterior-inferior LGN (Fig. 4A). Individual offsets corroborate this bias, with a posterior-inferior vector of 3.9 mm (3.4 mm posterior, 1.7 mm inferior). LGN retinotopy makes this bias systematic, where the upper visual field maps to inferior LGN, and central eccentricities to posterior LGN, with eccentricity increasing anteriorly^22,23^ (Fig. 4A). A posterior-inferior bias therefore preferentially modulates the upper visual field at central-to-parafoveal eccentricities.

**Fig. 4:**
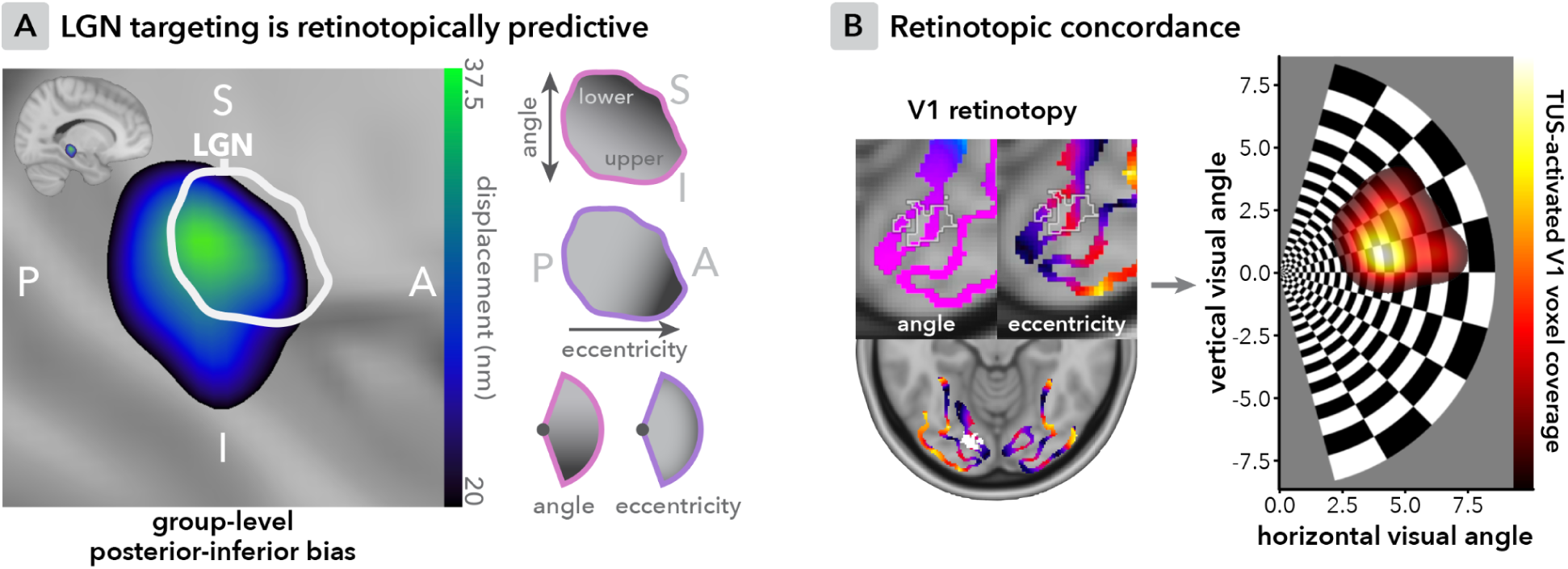
Retinotopic concordance between targeted region of LGN and activated region of V1. **(A)** Group-average MR-ARFI displacement across participants in standard space, revealing a posterior-inferior targeting bias that corresponds retinotopically to the upper visual field at central-to-parafoveal eccentricities. **(B)** Left: The Benson retinotopic atlas in standard space overlaid with the LGN-TUS-activated V1 cluster (outline ROI in top panel, white voxels in bottom panel). Right: The colocalized Benson retinotopy and LGN-TUS-activated V1 cluster were used to estimate the modulated retinotopic extent. The upper-right visual field was significantly engaged, aligning retinotopically with the LGN region of highest exposure.

To test this, we projected the TUS-activated V1 cluster onto the visual field using the Benson retinotopic atlas^24^ (Fig. 4B). The cluster reflects neuromodulation (LGN-TUS>Control-TUS), with the stimulus-driven checkerboard response subtracted. These voxels mapped directly onto the upper-right quadrant at a median eccentricity of 4.73° (IQR: 4.04°-5.38°) and polar angle of 75.81° (IQR: 65.89°-83.58°). These results provide additional converging evidence of a local neuromodulatory effect in LGN being expressed transsynaptically in downstream cortex.

### Behavioral effects

During the fMRI scans, participants concurrently performed a contrast decrement detection task, wherein participants reported whether they perceived a decrement in the upper or lower quadrant of the contrast-reversing hemifield checkerboard (Supplementary Note 1-2; Supplementary Fig. 5,7). We found no robust group-level effects of LGN-TUS on contrast decrement detection accuracy or reaction time. As the psychometric function is steepest near threshold, the behavioral measure was expected to be most sensitive to LGN-TUS when performance was held near 75% correct. However, only eight participants performed within the intended 70-80% range during the experiment, with the remainder performing above it, likely owing to practice effects following the threshold estimation. Detection accuracy in those participants may therefore have been comparatively insensitive to any perturbation. An exploratory post-hoc analysis in those subjects revealed a trend in which LGN-TUS reduced accuracy. We interpret this trend as a preliminary indication of potential behavioral translation that will require the use of a more sensitive behavioral approach in future work (see Supplementary Note 1 and Supplementary Fig. 5 for details).

## Discussion

In this study, we found that in vivo MR-ARFI-quantified displacement explained inter-individual variation in the functional magnitude of ultrasound neuromodulation. By integrating MR-ARFI and TUS-fMRI, we demonstrated increases in local and network-level activation that scaled with empirically quantified dose. Our results thereby establish in vivo displacement as a significant source of the variability that currently limits robust TUS functional outcomes in humans. This work provides a proof-of-principle that MR-ARFI can facilitate the precision deep-brain neuromodulation required for robust causal neuroscientific research and, ultimately, effective personalized clinical intervention.

LGN-TUS caused a spatially specific increase in task-related activation relative to sonication of an active control site, both locally and in the downstream visual cortex (Fig. 2). We controlled the peripheral confounds that can masquerade as neuromodulation, particularly in online designs^25–28^, with empirically confirmed blinding. Effects were restricted to the sonicated hemisphere, providing secondary evidence of specificity considering potential confounding by the absence of visual drive in the unsonicated hemisphere. Prior studies targeting LGN have relied on downstream readouts from the visual cortex^11,15–17^ and behavior^15,17^. Here, we additionally find activation at the deep-brain target itself, allowing dose and response to be related at the target and then traced to a connected network.

Indeed, the magnitude of increase in task-related activation scaled with focal displacement on a between-subject level. This relationship was present both locally and in the monosynaptically connected ipsilateral visual cortex. We also observed a within-subjects relationship, where voxels exposed to greater displacement demonstrated stronger neuromodulatory responses. In sum, we find convergent evidence that displacement predicts the magnitude of neuromodulatory response in humans.

These results extend MR-ARFI-quantified displacement-dependent LGN-TUS effects on sheep visual evoked potentials^11^ to human hemodynamic activation. Our effects further align with established relationships between ex-vivo optically imaged displacement and neural spiking^29^, and between ultrasound-imaged displacement and hemodynamic activation in rodents, including the within-subject cross-voxel effect^20^. More broadly, similar functions have been reported between free-water amplitude and hemodynamic activation in non-human primates^30^ and rodents^31^. Taken together, across model systems and measurement modalities, displacement emerges as a key component of effective ultrasound neuromodulation.

Our results indicate system-level functional engagement that is consistent with a neural effect propagating transsynaptically along the geniculostriate pathway to the cortex, rather than a purely non-neuronal effect (e.g., exclusively vascular). This inference is supported by three observations. First, TUS produced downstream engagement of the visual cortex, the magnitude of which scaled with the local effect in LGN. Second, the LGN appears to mediate the network-level dose-response function in the expected direction. Third, the LGN subregion with the highest group-level displacement represents the same visual subfield as the significant cluster of V1 activation, such that downstream engagement followed retinotopy.

These network-level findings align with, and expand upon, prior reports quantifying downstream effects of LGN-TUS^11,16,17^. The retinotopic concordance between exposed LGN and engaged V1 in particular supports TUS spatial specificity, while additionally offering a noninvasive neuromodulatory complement to the correlational evidence underpinning human geniculostriate retinotopy^22^. This paradigm could enable causal mapping of subcortical visual organization in future work through dense sampling across multiple LGN subregions, referenced to individualized rather than template retinotopy.

Displacement can be leveraged to explain, and ultimately help control, a meaningful degree of inter-individual variability in circuit-level neuromodulatory response. The value of MR-ARFI in this pursuit is underscored by the specificity of explained inter-individual variance to empirically measured displacement. Simulated in-situ intensity predicted neither neuromodulatory response nor displacement itself. This discrepancy may stem from the established uncertainty in simulated transcranial intensity estimates^10^, which represents a lower bound that does not yet account for hair or air at the coupling interface. Note, however, that a single set of acoustic assumptions and a single simulation tool was used in this study. Comprehensive benchmarking of simulations against MR-ARFI requires dedicated research. A parallel explanation for the discrepancy is that acoustic simulations estimate intensity while MR-ARFI estimates displacement. The latter encodes viscoelastic interactions, thus providing an absorption- and tissue-mechanics-weighted metric rather than acoustic exposure alone, which may track operative dose with higher fidelity.

Importantly, the divergence between simulation and measurement concerned amplitude rather than location. Focal positions broadly agreed, with discrepancies on the order of our fiducial reconstruction error. Accurate focal position estimation with uncertainty surrounding amplitude corroborates prior reports on simulation performance^10,32–34^, now with heightened ecological validity in an in vivo human transcranial context. Limitations of amplitude estimation may explain the inconsistent detection of between-subject dose-response functions when relying on simulated in-situ exposure^27,35,36^. The discrepancy between simulated and measured amplitude in our sample might additionally caution against the increasingly considered practice of titrating applied free-water intensity to achieve more robust neuromodulation, when based on current simulations alone^16,37,38^ (Supplementary Fig. 3).

Instead, future work could leverage MR-ARFI to prescribe individualized dose in vivo. This proposal mirrors recommendations for empirically informed individualized dosing over simulation-based approaches in isolation, for instance using attenuation estimated by through-transmission^39^. We wish to stress that MR-ARFI is not a prerequisite for accurate neuromodulation, particularly given that its implementation is practically constrained by requiring an MRI. Simulated amplitude inaccuracies could alternatively be mitigated by improving coupling uniformity^9^ and implementing designs that emphasize spatial targeting or within-subject dose manipulation that are inherently more robust to simulated amplitude uncertainties. Indeed, our data support the utility of simulations for planning focal location. MR-ARFI itself could constitute an empirical benchmark against which simulations can be validated and improved.

We did not observe robust effects on behavior, which may reflect limited methodological sensitivity rather than an absence of behavioral translation per se. Across non-invasive brain stimulation, behavioral effects depend strongly on baseline performance and ongoing brain state, potentially reducing sensitivity at the group level^40,41^. Two features of our design may have limited sensitivity. First, in line with prior work^16^, we consider the spatial circumscription of the activated V1 cluster. This region represented a portion of the visual field that largely did not overlap our behavioral cues. Tight circumscription may also limit sensitivity of cortical electrophysiological readouts in this paradigm^15^. Second, sensitivity to neuromodulation was likely limited by practice effects that shifted performance from threshold-level, where perturbation is more readily detected^42,43^. In future work, adaptive psychometric thresholding could maintain a sensitive regime while capturing TUS effects through threshold dynamics. Cue placement could additionally be broadly aligned with the exposed retinotopic subfield or vice versa, and continuous oculomotor measures would offer a criterion-free alternative to detection accuracy.

Several limitations bound our findings. We tested only one free-water intensity per participant. Future work could additionally manipulate applied intensity to further characterize within-subject dose-response functions. In addition, given our sample size, independent replication is warranted before displacement is used prescriptively. The acoustic exposure required for MR-ARFI is an inherent limitation of the technique itself. These short pulses could in principle induce neuromodulatory effects of their own, although this is unlikely to account for the present results, as ARFI TUS pulses were not delivered during visual drive and the subsequent counterbalanced blocked design absorbs any residual interactions. Nonetheless, ARFI sequence optimization should minimize pre-experimental dose, and any effects thereof could be quantified directly in dedicated experiments. Given the proximity of the optic radiation to the LGN, we also cannot exclude concurrent engagement of white matter, which TUS is increasingly recognised to modulate^38,44,45^. Both routes converge on the same downstream readout, but cannot be disentangled here as our field of view did not permit displacement quantification across the optic radiation. Future research could explicitly disambiguate their contributions with differentiated targeting, and should ensure that the optic radiation is not exposed in control conditions. Finally, the scale of displacement in the present study should not be read as prescriptive. Prior displacement imaging work has suggested requisite amplitudes that are an order of magnitude higher for neuronal activation, albeit in rodents and at rest^20^. The model used in Eq. 1 to estimate displacement assumes linear tissue motion with time of the TUS application, which is approximately true from simulation^14^. Any minimum effective displacement is likely also conditional on brain state, anatomy, and temporal pulsing parameters. Moreover, dose-response functions themselves need not be linear or monotonic.

Our targeting characterizes the performance of a first-generation, largely manual implementation. Initial focal steering was informed by the geometric constellation of transducer fiducials and target, then manually re-steered based on realtime MR-ARFI with variable signal quality, in an approach similar to that proposed for nonhuman primates^46^. Robust group-level effects were obtained despite the millimeter-scale targeting variability of this initial approach. One interpretation of robust effects surviving targeting variability is that, as previously acknowledged^9,20^, the field of effective engagement could extend beyond the intensity full-width half-maximum conventionally used to describe TUS resolution^9,20^. Nevertheless, future implementations could automate fiducial reconstruction, focus localization, motion correction, and re-steering, along with improving MR-ARFI signal quality^13^, which will increase MR-ARFI-based targeting precision.

Beyond targeting, measured in vivo displacement provides a path to investigate the primary biophysical driving forces underlying ultrasound neuromodulation. Displacement simultaneously encodes normal strain, shear strain, and acoustic particle displacement, which represent parallel, likely synergistic driving mechanisms of ultrasound neuromodulation. Normal and shear strain could be dissociated by varying applied amplitude and focal offset, while high bandwidth transducers could separate acoustic radiation force from particle displacement^8^. Our design affords neither the requisite combinations of amplitude and offset nor the MR-elastography such work demands. Nonetheless, MR-ARFI offers an accessible route to testing these manipulations in humans.

Our findings establish a framework for quantifying, explaining, and ultimately reducing variability in the local and network-level neuromodulatory effects of TUS through empirical quantification of in vivo displacement. LGN-TUS elicited a spatially specific increase in functional engagement both locally and in the downstream ipsilateral visual cortex, and MR-ARFI confirmed that the strength of this effect is indeed predicted by displacement. Our results provide a sensory analogue of the circuit-level intervention increasingly considered for understanding and treating clinical conditions^18,19^. More broadly, the functional magnitude of subcortical ultrasound neuromodulation can now be quantitatively linked to in vivo displacement, opening new opportunities for the controlled causal investigation of distributed human brain circuits.

## Methods

### Participants

Twenty-four healthy adults (mean age 28.3 ± 7.5 years, range 19-49; 14 male) completed the study. Participants were screened for contraindications to TUS and MRI, including for history of brain surgery or serious head trauma, current neuropsychiatric diagnoses, use of psychoactive medication, and the presence of ferromagnetic material in the body. All participants had normal or corrected-to-normal vision. Written informed consent was obtained from each participant in accordance with the Declaration of Helsinki, and the study was approved by the Stanford University Institutional Review Board.

### Visual Stimuli

Visual stimuli comprised single hemifield 8 Hz contrast reversing polar checkerboards, presented during MRI on a 60 Hz projector at a viewing distance of 33 cm (see Supplementary Fig. 6 for details). These stimuli served to evoke lateralized visual BOLD responses in the LGN and visual cortices to support LGN localization and to elicit an ongoing visual response that could be modulated by TUS (Fig. 1C).

As a secondary measure to fMRI, we assessed TUS effects on visual perception using a two-alternative contrast decrement localization task. Contrast decrements were superimposed on the contrast reversing polar checkerboards, applied to a patch in the upper or lower quadrant in pseudorandomized order. Participants were required to report the quadrant in which the decrement appeared. Accuracy and reaction time were recorded. The contrast decrement was 125 ms in duration, phase-locked to contrast reversal at fixed time points per block, and presented at the perceptual threshold level estimated by an adapted parameter estimation by sequential testing algorithm. Clearly visible (i.e., high contrast decrement) probes were interspersed to monitor compliance. Summed visual input was matched across all experimental conditions within each block and run, ensuring that differences in visual drive would not confound the primary BOLD signal comparisons. See Supplementary Note 2 and Supplementary Fig. 7 for details on the behavioral task.

### MRI acquisition and analysis

#### Structural scans

Imaging was performed at the Stanford Lucas Center for Imaging on a 3T GE SIGNA Premier scanner (GE Healthcare, WI, USA; 70 mT/m gradients, 170 T/m/s slew rate) with a 48-channel head coil. A higher-order shim was computed over the deep brain, including the LGN, to diminish spatial distortion and optimize signal-to-noise ratio. Together with the use of the 48-channel head coil, this may contribute to the successful detection of effects on LGN BOLD signal in this study compared to prior work^16^. Anatomical acquisitions comprised T1-weighted, T2-weighted, and zero-echo-time (ZTE) images were acquired. Functional data were acquired with single-shot spiral in-out gradient-echo readout^47^. Cardiac and respiratory activity were recorded for physiological nuisance correction at the post-processing stage. Individual native space images were aligned by normalized mutual information rigid transformations using FSL^48^, and statistical maps were warped to standard MNI152 space using ANTs^49^. Full sequence and physiological recording parameters are reported in Supplementary Table 1.

#### fMRI scans

A visual localizer scan was acquired in an oblique plane perpendicular to the axis of acoustic propagation for targeting and to permit subsequent MR-ARFI slice subselection (see *MR-ARFI*). The slice prescription covered the left LGN and visual cortices. Real-time processing for functionally informed targeting employed a GLM with physiological denoising using custom in-house C programs to provide lateralized visual activation maps. For the main experiment, two ten-minute neuromodulation fMRI scans were acquired, with axial-oblique slice coverage encompassing LGN and V1 in both hemispheres.

#### fMRI postprocessing and analysis

Functional data were analyzed in FSL FEAT with MCFLIRT motion correction, 3 mm FWHM smoothing, high-pass filtering (180 s for neuromodulation scans, 90 s for the localizer), FILM prewhitening, and improved brain masking based on a resliced SimNIBS v4.5 CHARM segmentation^50^. Physiological noise was modelled with FSL PNM^51^ (16 regressors; Supplementary Table 1). Each ten-minute neuromodulation run was modelled with four boxcar regressors (LGN-TUS, Control-TUS, and no-TUS Baseline during right- or left-hemifield stimulation), convolved with a double-gamma HRF and its temporal derivative. Per participant, neuromodulation scans were aggregated using FEAT. The primary contrast was the hypothesized LGN-TUS>Control-TUS effect.

The left (target) LGN ROI was defined from both functional and structural data, combining the localizer right>left visual contrast (z > 2.3) with anatomical LGN as delineated by HIPS-THOMAS thalamic statistical segmentation^52^. Given that the localizer covered only the left LGN, the right anatomical LGN segmentation was volume-matched to the left ROI so that left-right comparisons were not confounded by ROI size. For the small-volume LGN, statistical inference was based on two-tailed paired t-tests on estimates in the ROI. Downstream effects on the much larger visual cortices were tested by warping each participant’s LGN-TUS>Control-TUS estimate and its variance to MNI152 standard space and using FSL FLAME1+2 group-level mixed-effects analysis constrained to a bilateral V1 mask, mirroring the approach taken by Martin and colleagues (2025)^16^. The cluster forming threshold was z > 2.3 with cluster-corrected p < 0.05.

### Transcranial ultrasound stimulation

Ultrasound stimulation was delivered using a 64-element phased array transducer (Imasonic SAS, Voray-sur-l’Ognon, France) driven by an IGT generator (Image Guided Therapy, Pessac, France). Stimulation was applied in an online approach at a fundamental frequency of 500 kHz, a free-water spatial-peak pulse-average intensity (I_SPPA_) of 50 W/cm^2^, and a 145 ms continuous pulse with 7 ms linear ramping to mitigate auditory and somatosensory co-stimulation^9,28,53^ (Fig. 2B). This continuous pulse was time-locked to threshold-level contrast decrements. In each 15-second block, the first epoch (1/8 block) had no stimulation. TUS was delivered in 6/7 of the remaining epochs (13.125 s). This could be characterized as an average pulse repetition frequency of 0.46 Hz. All stimulation parameters fell within established safety guidelines^54^ (see Supplementary Table 2).

The 500 kHz fundamental frequency was chosen to provide a balance between spatial resolution, mitigation of somatosensory co-stimulation over the temporal window^28^, and transcranial transmission efficiency. We delivered continuous rather than pulsed TUS, following the longer continuous pulses used in the only prior human LGN-TUS fMRI study^16^ and in an online human visual evoked potential TUS study^36^.

Individualized acoustic simulations were run using k-Plan (supplier/support: Brainbox Ltd., Cardiff, UK), a graphical user interface for k-Wave^55^. Simulations were informed by in-scanner fiducial-reconstructed transducer location and steering settings (see Supplementary Note 3 for acoustic simulation details and Supplementary Table 1 for ZTE parameters).

### MR-ARFI

Tissue displacement induced by acoustic radiation force was measured with MR-ARFI using the single-shot spiral time-series method previously established for the human brain^14^, using a variable-density single-shot spiral-out spin-echo sequence^47,56^. Displacement was encoded by a pair of inverted bipolar motion-encoding gradients (MEGs; 70 mT/m, 15 ms total), with two 7 ms ultrasound pulses delivered during the second lobe of each bipolar gradient, offset by a 2 ms ultrasound-MEG delay to maximise the encoded phase^14,57^ (Supplementary Fig. 8).

Displacement maps were computed based on an established pipeline^14^. Briefly, the within-brain phase time series was corrected for cardiac and respiratory fluctuations by complex RETROICOR^58^ and fitted voxelwise with a general linear model that was time-frame-weighted by inverse spatial variance of phase, in which the 25-time-frame off/on structure was the regressor of interest and the slicewise spatial-mean brain phase was a nuisance regressor to correct for bulk motion. This yielded a phase shift due to tissue displacement of β radians at each voxel. Physical displacement Δ*x* following:

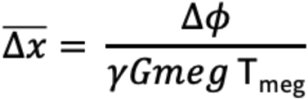

Where Δ**φ** is the phase difference, γ is the gyromagnetic ratio for protons, G_MEG_ (amplitude of motion-encoding gradients) = 70 mT/m, and T_MEG_ (duration of bipolar gradients) = 15 ms.

During MR-ARFI the free-water I_SPPA_ was 75 W/cm^2^, cycled in 25-frame off/on blocks over 100 frames. Three 4 mm-thick slices (1.72 × 1.72 mm in-plane) were selected from the oblique localizer prescription, centered on the localizer-identified LGN and perpendicular to the axis of acoustic propagation, resulting in a profile sampling 12 mm of the axial focus. MR-ARFI was acquired using a higher TUS intensity than applied for neuromodulation to increase sensitivity to individual differences. Given that radiation force scales linearly with intensity, and displacement with ARF at these amplitudes, the factor is constant across participants and leaves inferences unaffected.

### Experimental conditions

TUS was delivered to the left LGN and to the left temporal lobe grey matter as an active control site, which is not involved in early visual processing. This control site allowed us to separate direct neuromodulatory effects of LGN-TUS from nonspecific (peripheral) confounds^9,28^. During both LGN- and Control-TUS, the contrast reversing checkerboard was presented in the right visual hemifield. Baseline conditions involving no TUS were acquired with both right and left hemifield checkerboards, permitting cooling and enabling a control test for lateralized visual activation that confirmed task compliance and validated the analysis pipeline. These four conditions were presented in counterbalanced order across two ten-minute scans in 15-second pseudorandomized blocks (see Supplementary Fig. 9 for details). Within each block, sonication was delivered six times, time-locked to contrast decrement presentation, for a total of 870 ms per block (Fig. 2B).

### Masking & blinding

Throughout all conditions, Gaussian white noise was played continuously through pneumatic earbuds at each participant’s maximum tolerable volume. To further mitigate any audible differences between conditions, a salient 4 kHz masking tone overlapped the TUS interval during LGN-TUS and Control-TUS, or began at the equivalent time point during baseline (see Supplementary Fig. 7 for detailed experimental timing). The tone was presented with a leftward stereo bias to facilitate perceived origin from the transducer. Note that this tone provided additional auditory masking, rather than matching, as a confound-matched auditory stimulus was not achievable with the frequency response of the pneumatic audio system.

To quantify blinding, a two alternative forced choice detection task was administered at the end of the session, in-scanner, under the same conditions as the main experiment. LGN-TUS, Control-TUS, and right-hemifield Baseline were each presented eight times and participants reported whether they thought the stimulation was ‘real’. Sensitivity (d′) was estimated with log-linear correction^59^. Both TUS conditions were detectable relative to baseline (Supplementary Fig. 10). While this limits interpretability of any effects against baseline, unsuccessful blinding of baseline does not bear on the primary comparison of interest, namely, the difference between LGN-TUS and the active control condition. Indeed, participants could not discriminate LGN-TUS from Control-TUS (Fig. 2A). Moreover, sensitivity (d′) did not differ from zero, further confirming that the primary contrast was well blinded (mean d′ = −0.05; t(23) = −0.34, p = 0.735; 95% CI [−0.372, 0.266]; dz = 0.07; BF01 = 4.42).

### Procedure

The experiment was conducted in a single session. Hair preparation began approximately 30 minutes before the participant entered the scanner to increase hair soak time, and again immediately before coupling. Ultrasound gel (Aquasonic 100, Parker Laboratories) was applied over a broad area surrounding the temporal bone following ITRUSST practical guide recommendations^9^. Directly prior to scanning, additional gel was applied and the transducer was coupled to the head using a combination of gel and a 10° angled gel pad (Aquaflex, Parker Laboratories). The transducer was positioned anterosuperior to the left ear using a custom holder with a 10° posteroinferior tilt, which oriented the acoustic axis towards the LGN. The transducer was secured with a velcro elastic band and further stabilised by the MR head coil and cushions once the participant was supine.

During the initial structural scans, participants were trained on central fixation and on the behavioral contrast decrement detection task. Next, we acquired a five-minute localizer fMRI scan (300 volumes), in which hemifield checkerboard stimuli alternated between the left and right visual field. Contrast-decrement detection thresholds were estimated concurrently. This fMRI acquisition was analyzed on-scanner to delineate the LGN for targeting and, later, for independent ROI definition.

The acoustic beam was then guided to the localizer-defined LGN by phased electronic steering, leveraging the ±20 mm lateral and ±40 mm axial steering range afforded by the 64-element array and IGT generator. Initial steering settings were determined from the scanner coordinates of embedded fiducials on the transducer combined with the LGN location (Supplementary Fig. 4). MR-ARFI images were then acquired and processed in real time, and adjustments to the steering settings were made on the basis of these initial displacement estimates. At the final steering settings, MR-ARFI was used to quantify the in vivo displacement reported throughout.

Two ten-minute neuromodulation scans (600 volumes each) followed. Contrast decrements during these scans were presented at each participant’s estimated detection threshold. Blinding was assessed in the scanner at the end of the session.

### Quality control and exclusions

We incorporated two quality control checks. First, for the neuromodulation scans, we required left LGN to be more active during right than left visual stimulus presentation, thus confirming central fixation and task compliance. One run from a single subject was excluded where no such lateralisation was observed. For behavioral outcomes, data were not acquired for one participant due to technical issues. The second run of behavioral data was excluded for two participants with >25% missing trials (37.9% and 39.2%). In the remaining sample, participants responded on 99.6% of trials (IQR: 97.4-99.7%) and correctly identified high-contrast quality-control trials 93.8% of the time (IQR: 89.4-98.8%), demonstrating task compliance. Data required for MR-ARFI postprocessing were missing for the first three participants, and were therefore not included in analyses including displacement metrics. In summary, we analyzed N = 24 for primary fMRI analyses, N = 23 for behavioral analyses, and N = 21 for analyses including MR-ARFI metrics.

## Data analysis

To test local effects of LGN-TUS, we extracted ROI BOLD signal from the small, deep LGN and based statistical inference on paired t-tests. To test network-level effects on the larger visual cortex, we used second-level FSL FLAME1+2 analysis, mirroring prior work^16^. Hypothesized positive linear dose-response effects were confirmed with directional ordinary least squares (OLS) regression. The 95% confidence interval (CI) is reported throughout. Standardised effect sizes for within-participant comparisons are reported as Cohen’s d_z_, and regression effects are reported as standardised coefficients (β). Normality of paired differences and regression residuals was confirmed by visual inspection of distributions.

## Data availability

Data will be made available upon publication.

## Code availability

Code will be made available upon publication.

## Funding

This work was supported by the US National Institutes of Health through grants R21EY037048 (K.B.P., B.R.K., M.M., A.M.N., G.H.G.), R21EY037434 (K.B.P., B.R.K., M.M., A.M.N.), R01MH131684 (K.B.P., B.R.K., M.M., G.H.G.) and K08EY035037 (supporting R.T.A.). E. Fouragnan received support from a UK Research and Innovation Future Leaders Fellowship (MR/Y034368/1) and the Advanced Research and Invention Agency (SCNI-PR01-P15). R.T.A. additionally received support from AR-Bridge-To-Independence-00003163.

## Acknowledgements

We thank the Stanford Radiological Sciences Laboratory (RSL).

## Author contributions

Conceptualization: B.R.K., K.B.P., G.H.G., M.M., R.T.A; Data Curation: B.R.K., G.H.G.; Formal Analysis: B.R.K.; Funding Acquisition: K.B.P., R.T.A.; Investigation: B.R.K., M.M., E.F., M.T.W.S.; Methodology: B.R.K., K.B.P., G.H.G., M.M., E.F., R.T.A., M.T.W.S., A.M.N.; Project Administration: B.R.K., K.B.P.; Resources: K.B.P., G.H.G., A.M.N., R.T.A., B.R.K.; Software: B.R.K., G.H.G.; Supervision: K.B.P., E.F., E.F., G.H.G., R.T.A.; Visualization: B.R.K.; Writing -original draft: B.R.K.; Writing - review & editing: B.R.K., K.B.P., G.H.G., E.F., R.T.A., E.F., M.T.W.S., M.M., A.M.N.

## Competing Interests

B.R.K. is now employed by Attune Neurosciences. E. Fouragnan and K.B.P. serve as advisers to Attune Neurosciences. The remaining authors declare no competing interests.

## References

1. Murphy, K. & Fouragnan, E. The future of transcranial ultrasound as a precision brain interface. PLOS Biol. 22, e3002884 (2024).

2. Legon, W. et al. Transcranial focused ultrasound modulates the activity of primary somatosensory cortex in humans. Nat. Neurosci. 17, 322–329 (2014).

3. Meijer, S. et al. The human amygdala in threat learning and extinction. Sci. Adv. 10.1126/sciadv.aea8233 (2026) doi:10.1126/sciadv.aea8233.

4. Yaakub, S. N. et al. Non-invasive ultrasonic neuromodulation of the human nucleus accumbens impacts reward sensitivity. Nat. Commun. 16, 10192 (2025).

5. Clarke, S. et al. Multi-focal ultrasound neuromodulation to the dorsal anterior cingulate cortex disrupts behavioural and neural pain processing. Nat. Commun. 10.1038/s41467-026-72934-3 (2026) doi:10.1038/s41467-026-72934-3.

6. Pellow, C., Pichardo, S. & Pike, G. B. A systematic review of preclinical and clinical transcranial ultrasound neuromodulation and opportunities for functional connectomics. Brain Stimulat. 17, 734–751 (2024).

7. Nandi, T., Kop, B. R., Butts Pauly, K., Stagg, C. J. & Verhagen, L. The relationship between parameters and effects in transcranial ultrasonic stimulation. Brain Stimulat. 17, 1216–1228 (2024).

8. Nandi, T. et al. Biophysical effects and neuromodulatory dose of transcranial ultrasonic stimulation. Brain Stimulat. 18, 659–664 (2025).

9. Murphy, K. R. et al. A practical guide to transcranial ultrasonic stimulation from the IFCN-endorsed ITRUSST consortium. Clin. Neurophysiol. 10.1016/j.clinph.2025.01.004 (2025) doi:10.1016/j.clinph.2025.01.004.

10. Krokhmal, A., Simcock, I. C., Treeby, B. E. & Martin, E. A comparative study of experimental and simulated ultrasound beam propagation through cranial bones. Phys. Med. Biol. 70, 025007 (2025).

11. Mohammadjavadi, M. et al. Transcranial ultrasound neuromodulation of the thalamic visual pathway in a large animal model and the dose-response relationship with MR-ARFI. Sci. Rep. 12, 19588 (2022).

12. Phipps, M. A. et al. Practical targeting errors during optically tracked transcranial focused ultrasound using MR-ARFI and array-based steering. IEEE Trans. Biomed. Eng. 71, 2740–2748 (2024).

13. Sengupta, S., Phipps, M. A., Chen, L. M., Caskey, C. F. & Grissom, W. A. Alternating-contrast single-shot spiral MR-ARFI with model-based displacement map reconstruction. Magn. Reson. Med. 95, 457–464 (2026).

14. Mohammadjavadi, M., Ash, R. T., Glover, G. H. & Pauly, K. B. Optimization of MR acoustic radiation force imaging (MR-ARFI) for human transcranial focused ultrasound. Magn. Reson. Med. 94, 1060–1071 (2025).

15. Scott, M. T. et al. Effects of transcranial focused ultrasound stimulation to human lateral geniculate nucleus on visual perception and steady-state visual evoked potentials. 2026.07.30.741804 Preprint at 10.64898/2026.07.30.741804 (2026).

16. Martin, E. et al. Ultrasound system for precise neuromodulation of human deep brain circuits. Nat. Commun. 16, 8024 (2025).

17. Webb, T. D., Wilson, M. G., Odéen, H. & Kubanek, J. Sustained modulation of primate deep brain circuits with focused ultrasonic waves. Brain Stimulat. 16, 798–805 (2023).

18. Siddiqi, S. H., Kording, K. P., Parvizi, J. & Fox, M. D. Causal mapping of human brain function. Nat. Rev. Neurosci. 23, 361–375 (2022).

19. Horn, A. & Neumann, W.-J. From adaptive deep brain stimulation to adaptive circuit targeting. Nat. Rev. Neurol. 21, 556–566 (2025).

20. Kim, S., Kwon, N., Hossain, M. M., Bendig, J. & Konofagou, E. E. Displacement and functional ultrasound (fUS) imaging of displacement-guided focused ultrasound (FUS) neuromodulation in mice. NeuroImage 298, 120768 (2024).

21. Fritz, M. S., Kenny, D. A. & MacKinnon, D. P. The Combined Effects of Measurement Error and Omitting Confounders in the Single-Mediator Model. Multivar. Behav. Res. 51, 681–697 (2016).

22. Kastner, S. et al. Functional Imaging of the Human Lateral Geniculate Nucleus and Pulvinar. J. Neurophysiol. 91, 438–448 (2004).

23. Rahmati, M., Curtis, C. E. & Sreenivasan, K. K. Mnemonic representations in human lateral geniculate nucleus. Front. Behav. Neurosci. 17, (2023).

24. Benson, N. C. & Winawer, J. Bayesian analysis of retinotopic maps. eLife 7, e40224 (2018).

25. Guo, H. et al. Ultrasound Produces Extensive Brain Activation via a Cochlear Pathway. Neuron 98, 1020–1030.e4 (2018).

26. Sato, T., Shapiro, M. G. & Tsao, D. Y. Ultrasonic Neuromodulation Causes Widespread Cortical Activation via an Indirect Auditory Mechanism. Neuron 98, 1031–1041.e5 (2018).

27. Kop, B. R. et al. Auditory confounds can drive online effects of transcranial ultrasonic stimulation in humans. eLife 12, RP88762 (2024).

28. Kop, B. R., de Jong, L., Kim, B. P., den Ouden, H. E. M. & Verhagen, L. Parameter optimisation for mitigating somatosensory confounds during transcranial ultrasonic stimulation. Brain Stimulat. 18, 1224–1236 (2025).

29. Menz, M. D. et al. Radiation Force as a Physical Mechanism for Ultrasonic Neurostimulation of the *Ex Vivo* Retina. J. Neurosci. 39, 6251–6264 (2019).

30. Yang, P.-F. et al. Differential dose responses of transcranial focused ultrasound at brain regions indicate causal interactions. Brain Stimulat. 15, 1552–1564 (2022).

31. Yuan, Y., Wang, Z., Liu, M. & Shoham, S. Cortical hemodynamic responses induced by low-intensity transcranial ultrasound stimulation of mouse cortex. NeuroImage 211, 116597 (2020).

32. Angla, C., Larrat, B., Gennisson, J.-L. & Chatillon, S. Transcranial ultrasound simulations: A review. Med. Phys. 50, 1051–1072 (2023).

33. Aubry, J.-F. et al. Benchmark problems for transcranial ultrasound simulation: Intercomparison of compressional wave models. J. Acoust. Soc. Am. 152, 1003–1019 (2022).

34. Miscouridou, M., Pineda-Pardo, J. A., Stagg, C. J., Treeby, B. E. & Stanziola, A. Classical and Learned MR to Pseudo-CT Mappings for Accurate Transcranial Ultrasound Simulation. IEEE Trans. Ultrason. Ferroelectr. Freq. Control 69, 2896–2905 (2022).

35. Yaakub, S. N. et al. Transcranial focused ultrasound-mediated neurochemical and functional connectivity changes in deep cortical regions in humans. Nat. Commun. 14, 5318 (2023).

36. Dunsford, S., Murphy, K., Darrieutort, E., Fouragnan, E. & Ganis, G. Unilateral online ultrasound stimulation of early visual cortex suppresses responses to contralateral visual stimuli. Brain Stimulat. 19, 103023 (2026).

37. Kim, H.-C., Lee, W., Weisholtz, D. S. & Yoo, S.-S. Transcranial focused ultrasound stimulation of cortical and thalamic somatosensory areas in human. PLOS ONE 18, e0288654 (2023).

38. Bao, S., Kim, H., Shettigar, N. B., Li, Y. & Lei, Y. Personalized depth-specific neuromodulation of the human primary motor cortex via ultrasound. J. Physiol. 602, 933–948 (2024).

39. Riis, T., Feldman, D., Mickey, B. & Kubanek, J. Controlled noninvasive modulation of deep brain regions in humans. Commun. Eng. 3, 13 (2024).

40. Silvanto, J., Bona, S., Marelli, M. & Cattaneo, Z. On the Mechanisms of Transcranial Magnetic Stimulation (TMS): How Brain State and Baseline Performance Level Determine Behavioral Effects of TMS. Front. Psychol. 9, (2018).

41. Hartwigsen, G. & Silvanto, J. Noninvasive Brain Stimulation: Multiple Effects on Cognition. Neurosci. Rev. J. Bringing Neurobiol. Neurol. Psychiatry 29, 639–653 (2023).

42. Farboud, S. et al. Rapid modulation of choice behavior by ultrasound on the human frontal eye fields. Nat. Commun. 17, 2966 (2026).

43. Butler, C. R. et al. Transcranial ultrasound stimulation to human middle temporal complex improves visual motion detection and modulates electrophysiological responses. Brain Stimulat. 15, 1236–1245 (2022).

44. Tsuchiyagaito, A. et al. Reversible modulation of a deep white matter surgical target for depression with low-intensity focused ultrasound. Neuropsychopharmacology 51, 612–621 (2026).

45. Curtin, D., Walinjinkar, J., Fouragnan, E. F., Hendrikse, J. J. & Coxon, J. P. Transcranial ultrasound stimulation induces white matter plasticity in the human corticospinal tract. 2025.11.10.687582 Preprint at 10.1101/2025.11.10.687582 (2025).

46. Mishra, A. et al. Disrupting nociceptive information processing flow through transcranial focused ultrasound neuromodulation of thalamic nuclei. Brain Stimulat. 16, 1430–1444 (2023).

47. Chang, C. & Glover, G. H. Variable-density spiral-in/out functional magnetic resonance imaging. Magn. Reson. Med. 65, 1287–1296 (2011).

48. Jenkinson, M., Beckmann, C. F., Behrens, T. E. J., Woolrich, M. W. & Smith, S. M. FSL. NeuroImage 62, 782–790 (2012).

49. Avants, B. B. et al. A reproducible evaluation of ANTs similarity metric performance in brain image registration. NeuroImage 54, 2033–2044 (2011).

50. Puonti, O. et al. Accurate and robust whole-head segmentation from magnetic resonance images for individualized head modeling. NeuroImage 219, 117044 (2020).

51. Brooks, J. C. W. et al. Physiological noise modelling for spinal functional magnetic resonance imaging studies. NeuroImage 39, 680–692 (2008).

52. Saranathan, M. et al. Comprehensive Segmentation of Deep Grey Nuclei From Structural MRI Data. Hum. Brain Mapp. 46, e70350 (2025).

53. Mohammadjavadi, M. et al. Elimination of peripheral auditory pathway activation does not affect motor responses from ultrasound neuromodulation. Brain Stimulat. 12, 901–910 (2019).

54. Aubry, J.-F. et al. ITRUSST consensus on biophysical safety for transcranial ultrasound stimulation. Brain Stimulat. 18, 1896–1905 (2025).

55. Treeby, B. E. & Cox, B. T. k-Wave: MATLAB toolbox for the simulation and reconstruction of photoacoustic wave fields. J. Biomed. Opt. 15, 021314 (2010).

56. Glover, G. H. Spiral imaging in fMRI. NeuroImage 62, 706–712 (2012).

57. Kaye, E. A. & Pauly, K. B. Adapting MRI acoustic radiation force imaging for in vivo human brain focused ultrasound applications. Magn. Reson. Med. 69, 724–733 (2013).

58. Glover, G. H., Li, T.-Q. & Ress, D. Image-based method for retrospective correction of physiological motion effects in fMRI: RETROICOR. Magn. Reson. Med. 44, 162–167 (2000).

59. Hautus, M. J. Corrections for extreme proportions and their biasing effects on estimated values ofd′. Behav. Res. Methods Instrum. Comput. 27, 46–51 (1995).

