## Supplementary Material for "Dose-dependent ultrasound neuromodulation of the human brain"

#### Supplementary Figure 1 - MR-ARFI images

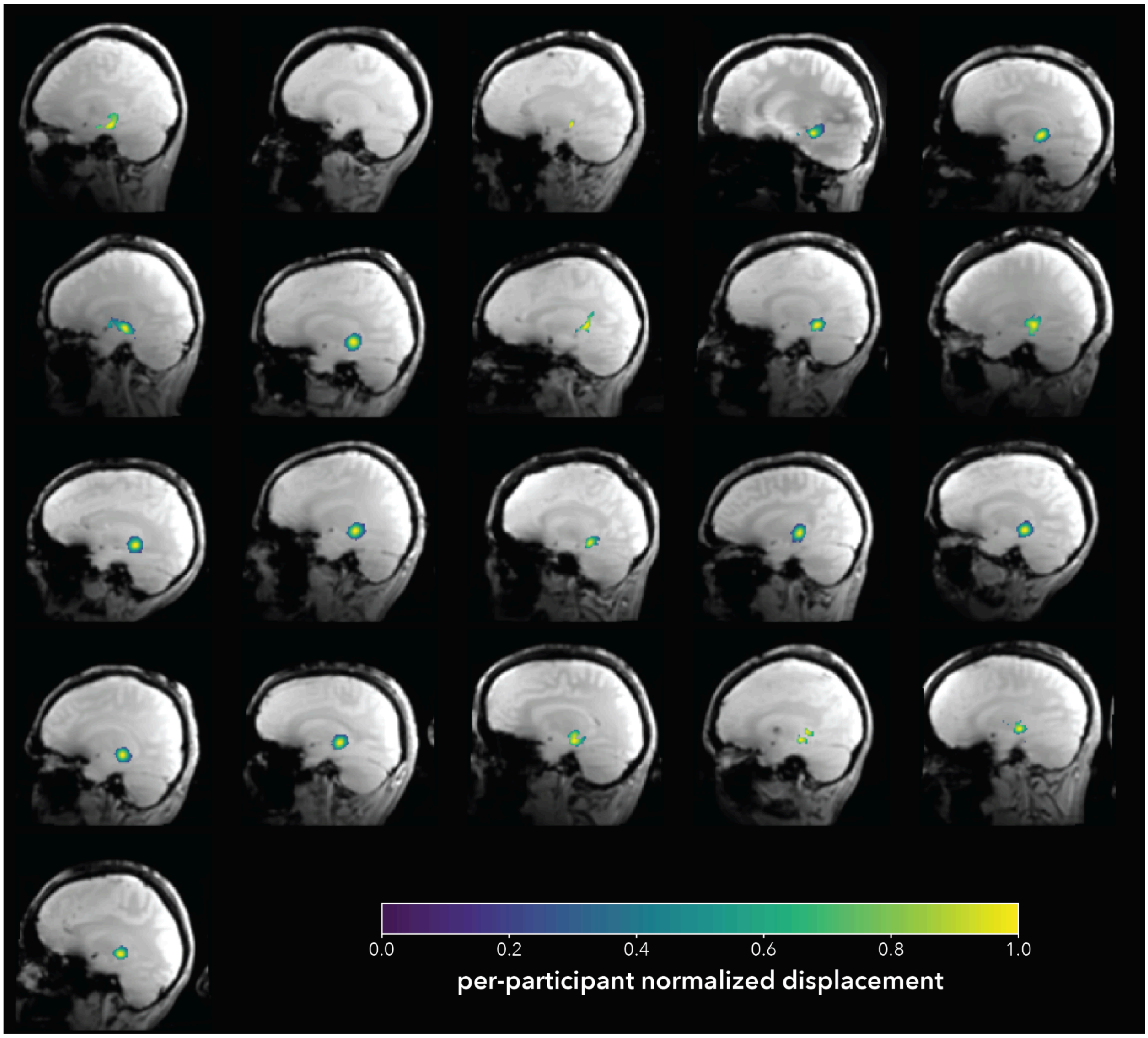


**Supplementary Fig. 1: MR-ARFI displacement images across 21 participants.** Participant-level peak normalized displacement maps overlaid on each participant's anatomical image (oblique). Light Gaussian smoothing is applied for visualisation only (σ = 1.5 voxels). Overlay voxels are restricted to significant displacement (i.e., t-score > 2.3). N = 21.

#### Supplementary Figure 2 - Mediation

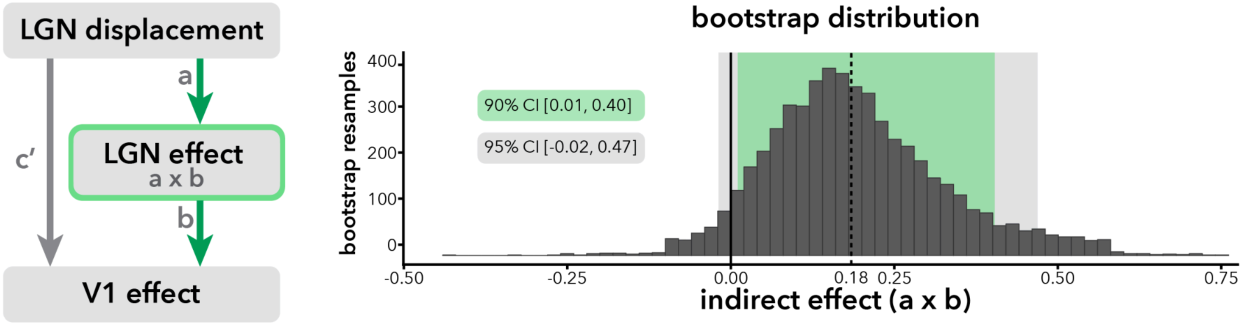


**Supplementary Fig. 2: LGN dose-response mediation.** Peak LGN displacement (X) → LGN effect (M) → V1 effect (Y); 5000 sample bootstrap. Both constituent paths were individually supported (a: β = 0.43, 95% CI [0.04, 0.90]; b: β = 0.43, 95% CI [0.01, 0.96]), while the direct path was not (c′: β = 0.19, 95% CI [−0.15, 0.68]). The indirect effect, i.e., the formal test of mediation, showed a positive trend (β = 0.18, 90% CI [0.01, 0.40]) but its 95% CI narrowly included zero ( 95% CI [−0.02, 0.47]). Bootstrap intervals are wide at this sample size, and measurement error in a mediator is known to attenuate the indirect path and inflate the direct path. We therefore treat this analysis as consistent with, but not formal evidence of, geniculostriate displacement-dependent propagation. N = 21.

**Supplementary Figure 3 - Acoustic simulations do not significantly predict displacement**

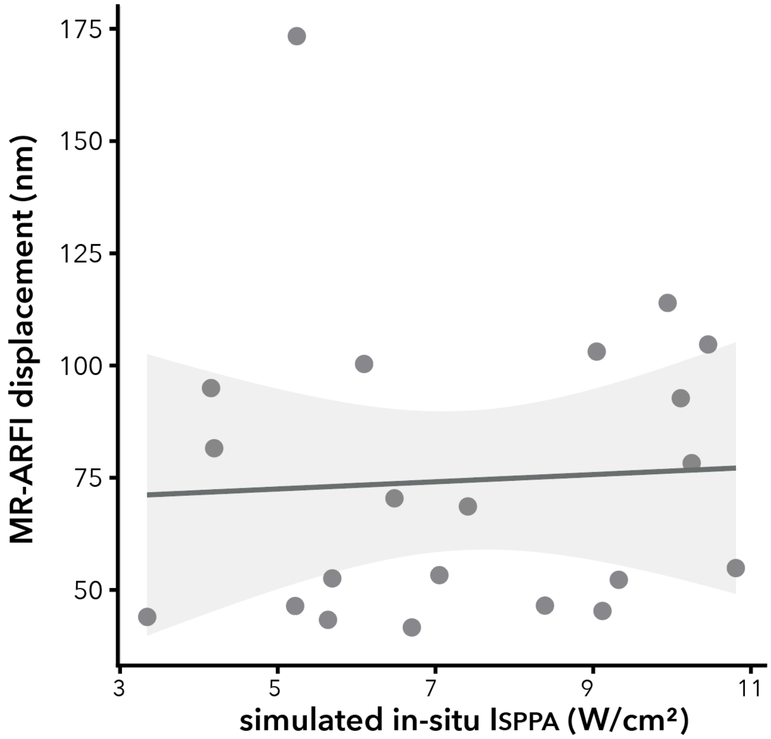


**Supplementary Fig. 3: Simulated in-situ intensity does not significantly predict MR-ARFI-quantified displacement.** Simulated in-situ intensity and measured in vivo displacement were uncorrelated (β = 0.13, 95% CI [-0.35, 0.61], t(19) = 0.57, p = 0.288 (one-sided), $R^2$ = 0.02). This discrepancy is unsurprising and likely reflects: (1) established uncertainty in simulated amplitude; (2) simulations not accounting for coupling quality, i.e., air at the coupling interface, whose impact is captured by MR-ARFI; and (3) the absence of elastography, which precludes converting measured displacement back to intensity in this sample. N = 21.

#### Supplementary Figure 4 - Transducer reconstruction

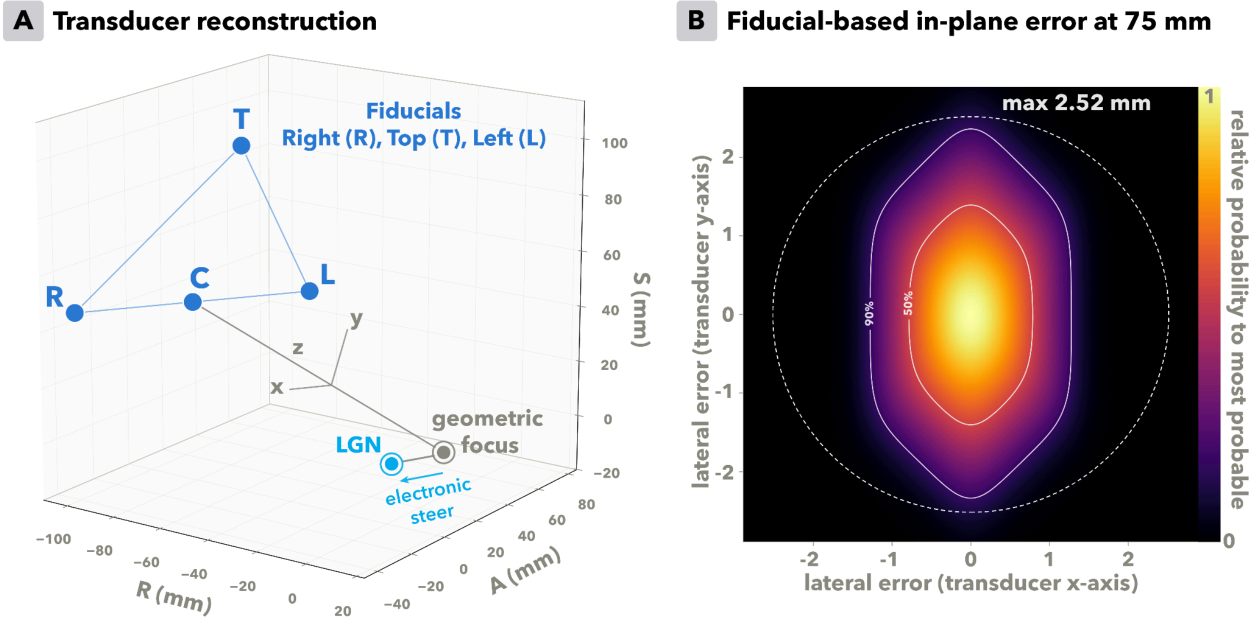


**Supplementary Fig. 4: Transducer reconstruction and fiducial error. (A)** Right-anterior-superior (RAS) coordinates of fiducials (dark blue) on the right (R), top (T), and left (L) of the transducers were used to reconstruct the center (C) and orientation of the transducer. The center of the fiducials was shifted -16 mm along *z* so the origin was at the back of the radiating surface. The geometric focus for our transducer was then identified at +75 mm along *z*. The RAS coordinate of LGN was used to determine the requisite initial electronic steering. **(B)** Fiducial coordinates were manually assigned by the researcher. The maximum plausible error per fiducial was 0.75 mm from the true fiducial center, which can introduce error in expected/simulated focal location, particularly at a ~75 mm focal depth. The probabilistic consequence of this error is depicted, illustrating that the in-plane focus cross section can deviate up to 2.52 mm depending on the combination of fiducial marking errors across the three fiducials. It is ~50% likely that the in-plane error is 1 mm or less.

#### Supplementary Note 1 - Behavioral Results

LGN-TUS did not significantly modulate accuracy or reaction times relative to Control-TUS at the group level (Supplementary Fig. 5A; accuracy: LGN-TUS = 82.5 ± 2.6%, Control-TUS = 83.6 ± 2.2% [M ± SE]; MΔ = −1.11 ± 0.97%; t(22) = −1.14, p = 0.268; 95% CI [−3.13, 0.91]; $d_z$ = −0.24; reaction time: LGN-TUS = 837 ± 17 ms, Control-TUS = 839 ± 18 ms; MΔ = −1.6 ± 3.8 ms; t(22) = −0.42, p = 0.678; 95% CI [−9.4, 6.2]; $d_z$ = −0.09).

This group-level null may reflect task insensitivity rather than an absence of behavioral translation per se. Our attempt to titrate decrements to individual threshold was initially successful but was not maintained (Supplementary Fig. 5D). Group accuracy over the first 10 trials was 76.1% (77.6% across the first 20), close to the intended 75%. Performance then drifted. Across the session as a whole, median accuracy across conditions was 82.4% (Supplementary Fig. 5B; IQR: 74.2–93.5%; M ± SD = 82.5 ± 11.0%; range 65.5–99.4%). This pattern is consistent with practice effects, and possibly fatigue later in the session, rather than indicating a miscalibrated starting threshold. We sampled a single decrement level, a deliberate tradeoff favouring more data at one level over noisier estimates across several. In the future, adaptive psychometric thresholding would better account for drift, but in the present study would have confounded visual input between condition blocks, introducing interpretability problems in our experiment using fMRI BOLD as the primary outcome.

Our data support the relevance of threshold-level baseline performance. Regressing the LGN-TUS versus Control-TUS accuracy difference on baseline accuracy revealed a significant positive relationship (Supplementary Fig. 5C; β = 0.184, SE = 0.080; t(21) = 2.29, p = 0.033; 95% CI [0.017, 0.351]; $R^2$ = 0.200), wherein participants with lower baseline accuracy showed stronger TUS-induced decreases. Eight participants remained within the intended range for baseline (Supplementary Fig. 5E). An exploratory analysis restricted to these subjects showed a trend that accuracy decreased during LGN-TUS (Supplementary Fig. 5E; LGN-TUS = 75.2 ± 2.5%, Control-TUS = 77.8 ± 2.9%; MΔ = −2.68 ± 1.21%; t(7) = −2.21, p = 0.063; 95% CI [−5.54, 0.19]; dz = −0.78), but with no corresponding effect on reaction time (LGN-TUS = 881 ± 20 ms, Control-TUS = 882 ± 20 ms; MΔ = −1.1 ± 5.5 ms; t(7) = −0.21, p = 0.840; 95% CI [−14.1, 11.8]; dz = −0.07). The direction of this effect is consistent with LGN-TUS increasing contrast gain. With the background at 100% Michelson contrast, a gain increase likely cannot raise the high contrast background to the same degree as the lower contrast decrement, thus compressing the perceived differential and reducing accuracy. However, this subsample is small and statistical evidence is weak-to-absent, thus requiring further research.

A parallel potential source of experimental insensitivity is that the significant V1 cluster did not retinotopically overlap either cue location (Supplementary Fig. 5F), so the neurally engaged region may not have been behaviorally relevant to this task. It remains possible, of course, that the neural effects simply did not translate to behavior.

#### Supplementary Figure 5 - Behavioral results

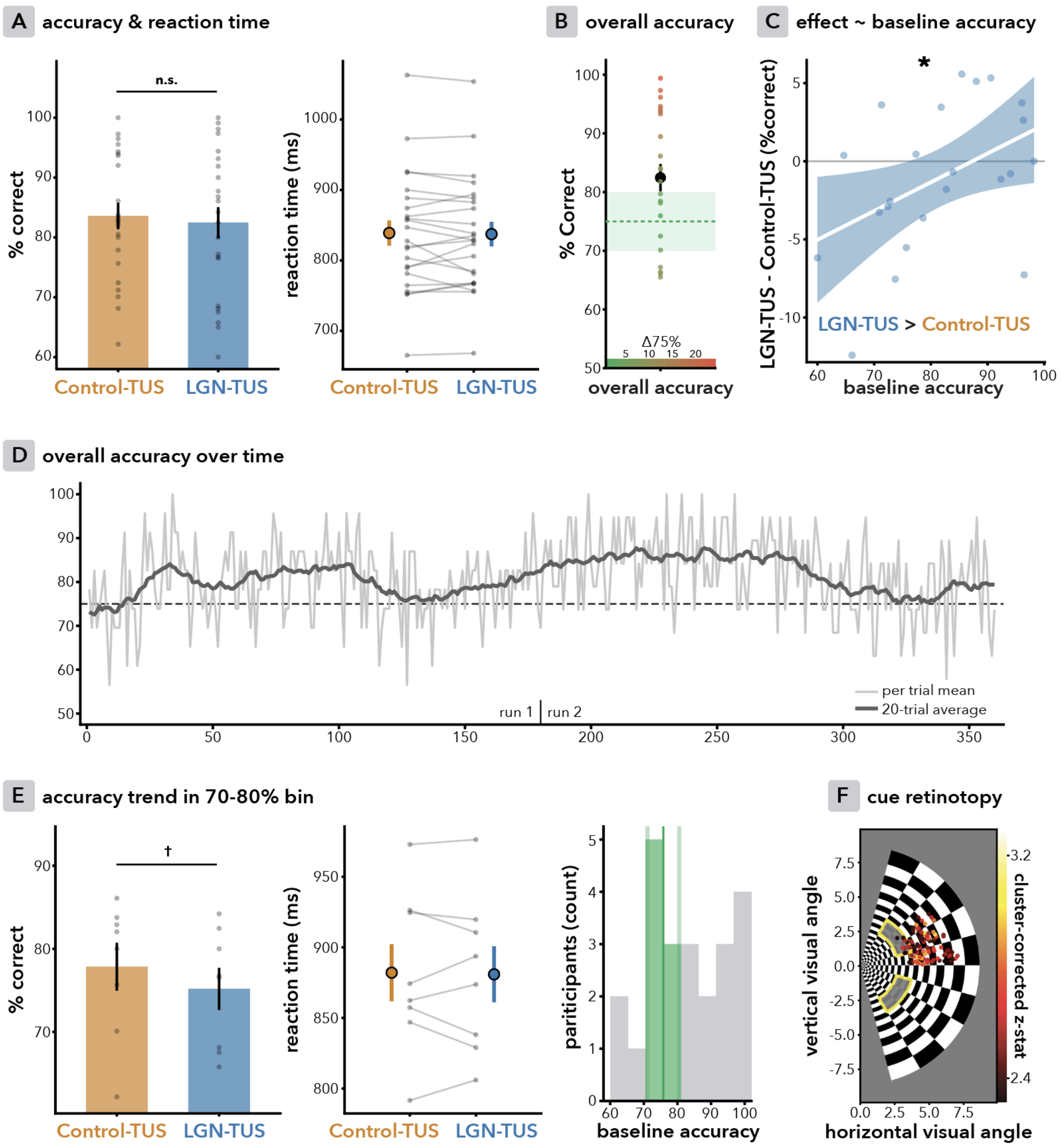


**Supplementary Fig. 5: Behavioral results. (A)** There was no significant group-level effect of TUS on contrast-decrement detection accuracy (left) or reaction time (right). **(B)** Although there were no ceiling or floor effects, overall accuracy exceeded the intended 75%, with most participants falling outside the 70–80% range. **(C)** Baseline accuracy significantly predicted the TUS effect, with lower baseline accuracy associated with stronger decreases in accuracy during LGN-TUS. **(D)** Group-level accuracy temporal dynamics likely reflect task practice effects, wherein participants start the experiment performing near threshold, but drift away from this threshold over time. **(E)** An exploratory analysis in the subset of eight participants whose baseline accuracy fell within the intended sensitive range showed a trend that accuracy was reduced during LGN-TUS as compared to active control (see Supplementary Note 1 for discussion). However, there was no effect on reaction time, and the sample is small. **(F)** A parallel potential explanation for the null is that the functionally engaged visual field, derived from the group-level significant V1 cluster of activation (see main text), did not overlap with either contrast decrement used in the behavioral task. N = 23.

#### Supplementary Figure 6 - Visual stimuli

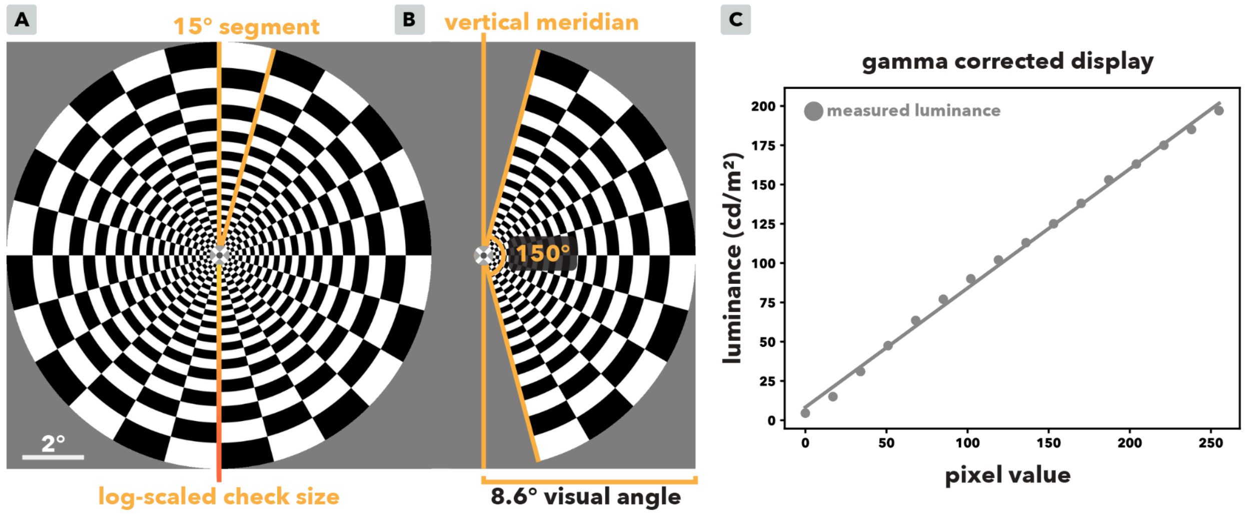


**Supplementary Fig. 6: Log-scaled polar checkerboard stimulus. (A)** The polar checkerboard stimulus was presented using a 60 Hz projector and subtended the central ~17.2° of the visual field (10 cm physical diameter at a 33 cm viewing distance) and was divided into 24 angular segments of 15° each (red lines), following prior work[1,2]. It comprised 28 concentric rings with a mean spatial frequency of 2 cycles/degree. Radial elements were log-scaled as a function of eccentricity to approximate the nonlinear mapping from the visual field to V1[3] and thus facilitate equal cortical representation across eccentricities[4]. Full (100%) Michelson contrast was used to maximise LGN drive[1], presented against a mid-grey background at a mean luminance of 105 cd/m$^2$. Participants fixated on a central bullseye fixation cross subtending a 0.6° visual angle. **(B)** On each block a single hemifield wedge was shown subtending 8.6° of the visual field. The wedge spanned 150° of polar angle, leaving a 15° buffer at the vertical meridian to avoid stimulating the contralateral hemifield. The checkerboard reversed contrast every 125 ms (i.e., 8 Hz reversal rate, 4 Hz temporal frequency), following established fMRI protocols for the LGN[1,5–8]. **(C)** Display luminance was measured with a MAVO-SPOT 2 photometer, and gamma correction was applied in PsychoPy to linearize the mapping between greyscale pixel values and luminance.

#### Supplementary Note 2 - Behavioral task

As a secondary measure, we assessed TUS effects on visual perception using a two alternative contrast decrement localisation task. Decrements were applied to a patch in the upper or lower quadrant of the concurrently 100% contrast-reversing hemifield (Supplementary Fig. 7). Participants reported which quadrant contained the decrement by button press and were instructed to guess if uncertain.

##### *Training*

During the initial structural scans (T1w, T2w, and ZTE), participants were trained to maintain fixation, to respond on every trial, and to respond correctly when the decrement was clearly visible. Training was repeated until these criteria were met. Trials without a response during neuromodulation scans were excluded from behavioral analyses (see main text for response rate).

##### *Contrast decrement perceptual threshold estimation*

During the localizer fMRI scan, each participant's psychometric function for contrast-decrement detection was estimated to avoid floor and ceiling effects, using an adapted parameter-estimation-by-sequential-testing (PEST) procedure (Supplementary Fig. 7). In sweep 1, a log-scaled range from 80% to 8% contrast decrement was presented over three blocks per hemifield. At the start of sweep 2, a logistic function was fitted to the sweep 1 data to provide an initial estimate of the 75% correct point (corresponding to 50% detection above chance for this two-alternative task). Sweep 2 then sampled seven contrast levels centered on this estimate, spanning ±75% of its value, over two blocks per hemifield. In the final sweep, the logistic function was iteratively refitted after every trial using all prior data from the relevant hemifield, and the next decrement was presented at the updated threshold, allowing the estimate to converge over the remaining five blocks per hemifield.

##### *Experimental timing during neuromodulation scans*

Each 15-second block was divided into eight 1.875 s epochs (Supplementary Fig 7). The first epoch of every block presented no decrement to absorb the elevated difficulty that pilot participants experienced on the first epoch following a hemifield switch. In six of the remaining seven epochs, contrast decrements were presented at the estimated perceptual threshold. These decrements appeared in the upper or lower quadrant in pseudorandomized order, counterbalanced across quadrants within each scan, within each block, and at each epoch position across blocks. In LGN- and Control-TUS conditions, ultrasound was delivered concurrently to the decrement, with an onset 20 ms prior. The single remaining epoch presented a clearly visible (high-contrast) decrement. Across blocks, this high-contrast epoch was interspersed as a compliance probe, pseudorandomized such that each epoch position contained at least one probe, with upper/lower presentations balanced. TUS was not administered during high-contrast decrements. Summed visual input was matched across the

LGN-TUS, Control-TUS, and Baseline conditions within each block and scan, ensuring that differences in visual drive could not confound the neuromodulatory contrast.

Contrast decrements were presented for 125 ms, phase-locked to the third contrast reversal in a given epoch (Supplementary Fig. 7). For LGN- and Control-TUS, a 145 ms sonication began 20 ms prior to the decrement. At the same time, for all conditions, a 4 kHz tone played over pneumatic earbuds. Throughout, white noise was played.

#### Supplementary Figure 7 - Behavioral task

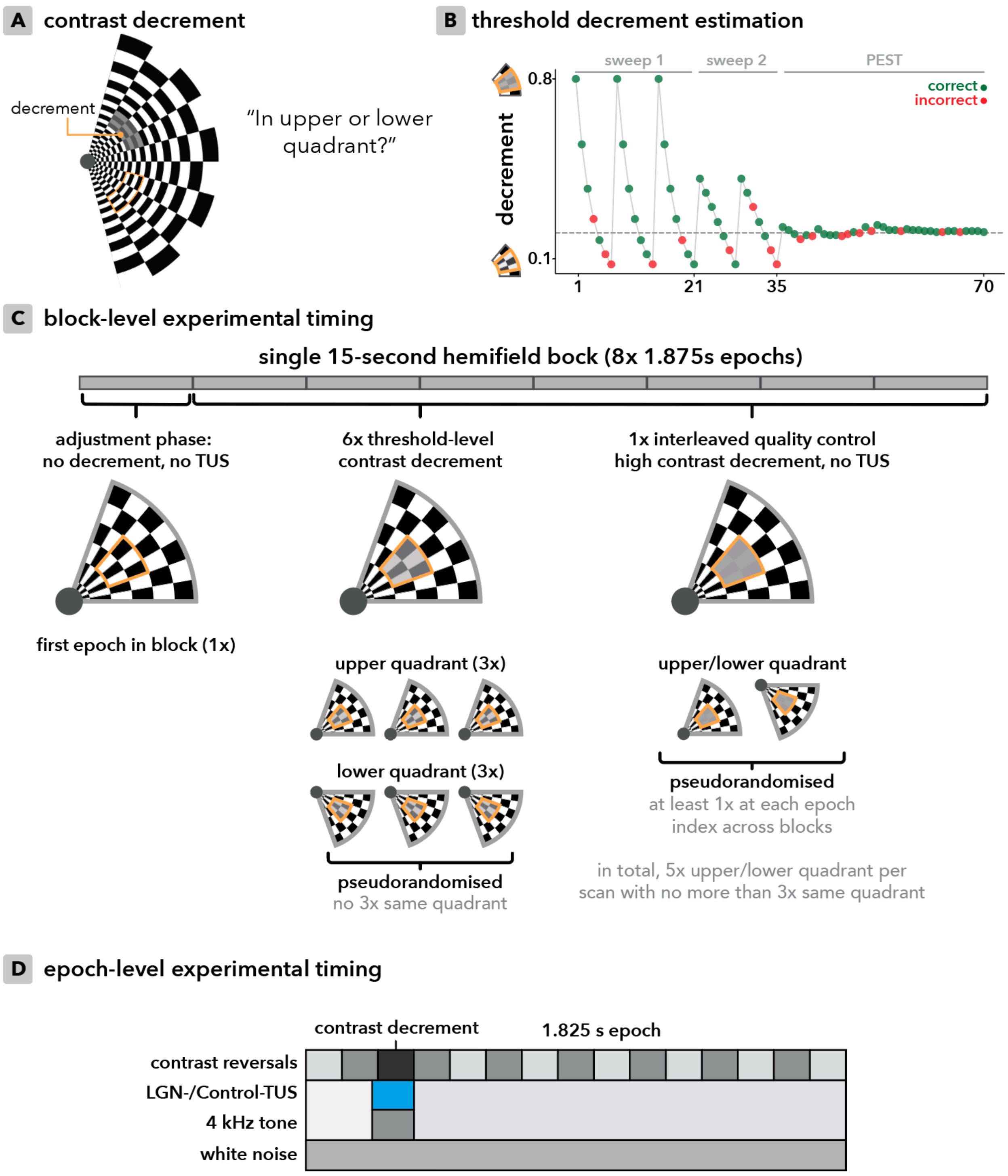


**Supplementary Fig. 7: Behavioral task design. (A)** A contrast-decrement was applied to a patch of the concurrently contrast-reversing hemifield radial checkerboard. Participants were required to respond whether they perceived the decrement in the upper or lower quadrant. **(B)**

An adapted parameter-estimation-by-sequential-testing (PEST) protocol was used to estimate contrast decrement perceptual thresholds, with two initial sweeps to inform the subsequent iterative logistic model threshold estimation and resampling. **(C)** Each 15-second block was subdivided into eight equal epochs, the first of which presented no contrast decrement, six of which presented the estimated threshold-level decrement, and one of which presented a high-contrast quality control decrement to monitor compliance. These conditions were counterbalanced such that visual drive was equal across experimental conditions. **(D)** Each contrast decrement lasted 125 ms, phase-locked to a single background contrast reversal. During LGN- and Control-TUS conditions, sonication began 20 ms prior and overlapped the decrement 125 ms decrement (i.e., 145 ms total). A 4 kHz tone was played over pneumatic earbuds at the same time, also in Baseline conditions. White noise played throughout.

### Supplementary Table 1 - MRI acquisition parameters

##### *Structural MRI acquisition parameters*

| Scan | TR (ms) | TE (ms) | Flip angle (°) | Voxel size (mm) | Acq. matrix | FOV (mm) | Slices | Bandwidth (Hz/px) |
|---|---|---|---|---|---|---|---|---|
| **T1w** | 5.556 | 2.192 | 8 | 0.508 × 0.508 × 0.8 | 300 × 256 | 260, 85% phase FOV | 220 | 244.1 |
| **T2w** | 2509 | 123.2 | 90 | 0.508 × 0.508 × 0.8 | 300 × 256 | 260, 85% phase FOV | 220 | 244.1 |
| **ZTE** | 37 | 0.016 | 1 | 0.469 × 0.469 × 0.8 | 300 × 300 | 240 | 220 | 244.1 |

##### *Functional MRI and MR-ARFI acquisition parameters*

| Scan | Type | TR (ms) | TE (ms) | Flip angle (°) | Voxel size (mm) | Acq. matrix | FOV (mm) | Slices | Frames |
|---|---|---|---|---|---|---|---|---|---|
| **Localizer** | SSFSE | 1000 | 30 | 62 | 1.72 × 1.72 × 4.0 | 80 × 80 | 220 | 16, oblique, perpendicular to axis of acoustic propagation, covering left LGN and V1 | 300 (+6 discarded) |
| **Neuromodulation runs** | GRE, Spiral-in/out | 1000 | 30 | 62 | 1.72 × 1.72 × 4.0 | 80 × 80 | 220 | 16, oblique, roughly stacked along axial plane, covering bilateral LGN and V1 | 600 (+6 discarded) |
| **MR-ARFI** | SE, Spiral | 800 | 50 | 90 | 1.72 × 1.72 × 4.0 | 80 × 80 | 220 | Subset of 3 slices from the localiser, centered on the LGN identified by the localiser | 100 (+6 discarded) |

##### *Physiological measurements and nuisance correction*

Cardiac and respiratory activity was recorded throughout with a finger photoplethysmograph (100 Hz) and a respiration belt (25 Hz) for physiological nuisance correction. Physiological regressors were generated with FSL[9] PNM from the recorded traces, including second-order cardiac, third-order respiratory, their interaction, respiration-volume-per-time, and heart rate (16 voxelwise regressors).

#### Supplementary Table 2 - TUS safety indices

| **Protocol** | **$MI_{TC}$** | **$TR_{TARGET}$** | **$TR_{SKULL}$** | **$TD_{TARGET}$** | **$TD_{SKULL}$** |
|---|---|---|---|---|---|
| neuromodulation | 0.87 | 0.17 °C | 1.42 °C | <0.1 CEM43°C | <0.1 CEM43°C |
| MR-ARFI | 1.06 | 0.11 °C | 0.82 °C | <0.1 CEM43°C | <0.1 CEM43°C |

*All metrics represent the maximum simulated metric, as this is the relevant benchmark for safety. Values are based on k-Plan simulation, and all values fall within established guidelines for biophysical safety of transcranial ultrasound stimulation[10].

#### Supplementary Note 3: Acoustic Simulations

Individualized simulations of acoustic wave propagation were performed using k-Plan, a graphical user interface for the pseudo-spectral time-domain solver k-Wave[11].

##### *Pseudo-CT generation*

Pseudo-CT (pCT) scans were generated to assign acoustic properties to the media (i.e., skull, soft tissue, and background). Specifically, zero-echo-time (ZTE) MRI scans were converted pCT using the 'petra-to-pct' toolbox[12]. First, the ZTE was debiased using the N4ITK MRI bias-field correction algorithm. Next, optimized custom tissue masks were designated, based on head tissue segmentation with the SimNIBS v4.5 CHARM pipeline using a custom atlas designed specifically to improve MRI-derived skull segmentation accuracy relative to CT ground truth for TUS[13]. Skull-mask voxels were then assigned Hounsfield Units via a linear mapping.

##### *Transducer reconstruction*

The 64-element Imasonic transducer was simulated in k-Plan at the highest available quality (per 2025, 'level 3/3'), designed to accurately simulate 2D and 3D pressure field hydrophone measurements repeated at several focal distances across the steering range. In absence of optical neuronavigation, in-scanner transducer position was derived from the right-anterior-superior (RAS) coordinates of three MRI-visible fiducials on the top, left, and right of the transducer, captured during the ZTE scan (Supplementary Fig. 4). From these coordinates, the transducer origin was reconstructed as the circumcenter of the left–top–right triangle, with an orientation vector ($z$) pointing toward the brain. The origin was translated −16 mm along $z$ to locate the back of the transducer's radiating surface, given that the fiducials sat 16 mm farther along $z$ relative to the back of the bowl. This reconstructed transducer position and orientation was defined with a 4×4 transformation matrix and embedded in a synthetic Localite instrument marker XML file, generated for compatibility with k-Plan's transducer position import function. Reconstructed transducer location was then visually confirmed in the graphical interface, and was indeed precisely aligned with the transducer in all cases, made visible by gel on the T1w image.

To assess robustness of this reconstruction approach, we determined that the realistic error between the true fiducial center and the researcher-marked center could deviate by maximally |0.75| mm. This seemingly minor error in transducer reconstruction can result in a meaningful shift of the focus at a ~75 mm focal depth. Therefore, we estimated the impact that fiducial marking error would have on the lateral cross-sectional focus location for every combination of plausible marking error across the fiducials. The potential error averaged about 1 mm, and reached up to 2.5 mm in the worst case (Supplementary Fig. 4). This error is equal to, or better than, the error one may expect for standard optical neuronavigation systems[14].

Nonetheless, this source of error should be considered in the comparison between ARFI and simulation peak location (main text Fig. 3C).

***Target reconstruction***

Simulating a given phased-array steering setting in k-Plan required the target to be defined in RAS+ coordinates, rather than as a steering setting on the transducer. We derived this coordinate by: (1) de-obliquing the pCT scan using AFNI's 3dWarp de-oblique function, which is required for coordinate correspondence when processing scans in k-Plan (2) using the reconstructed transducer, its geometric focus, and its applied steering settings along the transducer axes to derive the voxel corresponding to the administered steering on the de-obliqued scan, (3) converting the resulting coordinates from RAS to RAS+, and (4) setting the resulting coordinates as the target in the k-Plan interface. This procedure was performed independently for the LGN- and Control-TUS steering settings.

***Acoustic simulations in k-Plan***

A set of post-hoc simulations was run for LGN- and Control-TUS, with aberration correction and intensity rescaling turned off. With current k-Plan functionality (2025), this means that the amplitude of each element to 1, and elements are phased such that they would steer to the desired target in free-water. Simulations were run with identical transducer position and steering settings twice: once in free-water, and once with biological tissue acoustic properties assigned to the relevant media. The difference between these simulations thus captures the impact of biological tissue, most importantly the skull, on acoustic propagation. Free-water simulations had to be run for each individual subject, due to small differences caused by the transducer orientation and/or grid size that would otherwise confound the comparison between free-water and tissue-based outputs.

Output pressure fields were exported and re-aligned to native scanner space using the simulation grid LPI coordinates. Pressure amplitude was then converted to acoustic intensity as $I = P^2 / (2Z)$, with acoustic impedance $Z = \rho c$ computed for brain ($c = 1550$ m s$^{-1}$, $\rho = 1045$ kg m$^{-3}$). These initial pressures and intensities were then re-scaled by free-field calibration. Specifically, the free-water simulation was rescaled such that the spatial-peak pulse-average intensity ($I_{SPPA}$) matched that of the delivered protocol free-water intensity (i.e., 50 W/cm$^2$ for the neuromodulation protocol, 75 W/cm$^2$ for the ARFI protocol). The full transcranial model was then multiplied by this participant-specific scaling factor, yielding calibrated in-situ pressure and intensity maps. Finally, the transcranial mechanical index was then computed as MI = P (MPa) / $\sqrt{(f_0)}$, with $f_0 = 0.5$ MHz. Our LGN-TUS simulations show a maximum in-situ intensity of $7.98 \pm 0.49$ W/cm$^2$ (mean ± SE) at a 50 W/cm$^2$ free-water intensity, corresponding to ~16% transcranial transmission, which is commensurate with expected transmission through the thin temporal bone[15], aligned with previous reports[16,17], and thus interpreted as supporting evidence that the pipeline described above was appropriate.

##### *Simulated in-situ intensity and targeting*

The simulated maximum $I_{SPPA}$ in the LGN was 5.45 ± 0.57 (mean ± SE) W/cm$^2$ during LGN-TUS (Supplementary Fig. 11). During Control-TUS, the $I_{SPPA}$ in LGN was 0.37 ± 0.05 W/cm$^2$, confirming that we did not engage the LGN during the Control-TUS condition targeted at the grey matter of the temporal lobe (LGN-TUS exposure is ~15x stronger).

It is also pertinent to consider exposure of the optic radiation (OR), as white matter stimulation could plausibly contribute to the observed effects. Indeed, the left optic radiation was meaningfully exposed during LGN-TUS (6.66 ± 0.53 W/cm$^2$). For Control-TUS, exposure to the OR was 1.27 ± 0.22 W/cm$^2$ (LGN-TUS OR exposure is ~5x stronger). Here, the OR was defined as the binarized 50% probability thresholded Juelich histological atlas OR, warped to native space. We cannot exclude that exposure of the OR during LGN-TUS may contribute to the neuromodulatory effects reported here.

##### *Comparison between simulations and MR-ARFI-quantified displacement*

A rigorous quantitative comparison of simulated acoustic fields against MR-ARFI measurements is beyond the scope of this study and is best addressed by future work designed for that purpose. The present investigation was not optimized for such a comparison: (1) we sampled a three-slice volume (12 mm axial coverage) rather than the full beam, and so did not capture the entire axially elongated focus (≥ 25 mm); (2) we relied on a single simulation tool and a single set of acoustic-property assumptions, which need not match the estimates produced by alternative tools and assumptions; and (3) we did not acquire the MR-elastography data required to derive intensity from displacement, and therefore, unquantified subject- and region-specific viscoelastic properties prevent us from disambiguating intensity from displacement. An ideal future comparison would collect MR-elastography data, quantify the full acoustic beam with MR-ARFI rather than a single cross-section, and cross-validate across multiple acoustic assumptions and simulation packages.

Nonetheless, even with these limitations, simulations and MR-ARFI broadly correspond in peak location at the group level. For the group-level average displacement and intensity maps in standard space, the in-plane peak-displacement-to-peak-intensity distance differed by only 2 mm (main text Fig. 3C). We base this comparison on the group-average images so that subjects with higher displacement and a better-defined displacement peak naturally outweigh those with lower-quality MR-ARFI data, who would otherwise inflate the apparent simulation-to-ARFI discrepancy through a poorly constrained mix of measurement and simulation uncertainty.

#### Supplementary Figure 8 - MR-ARFI pulse sequence

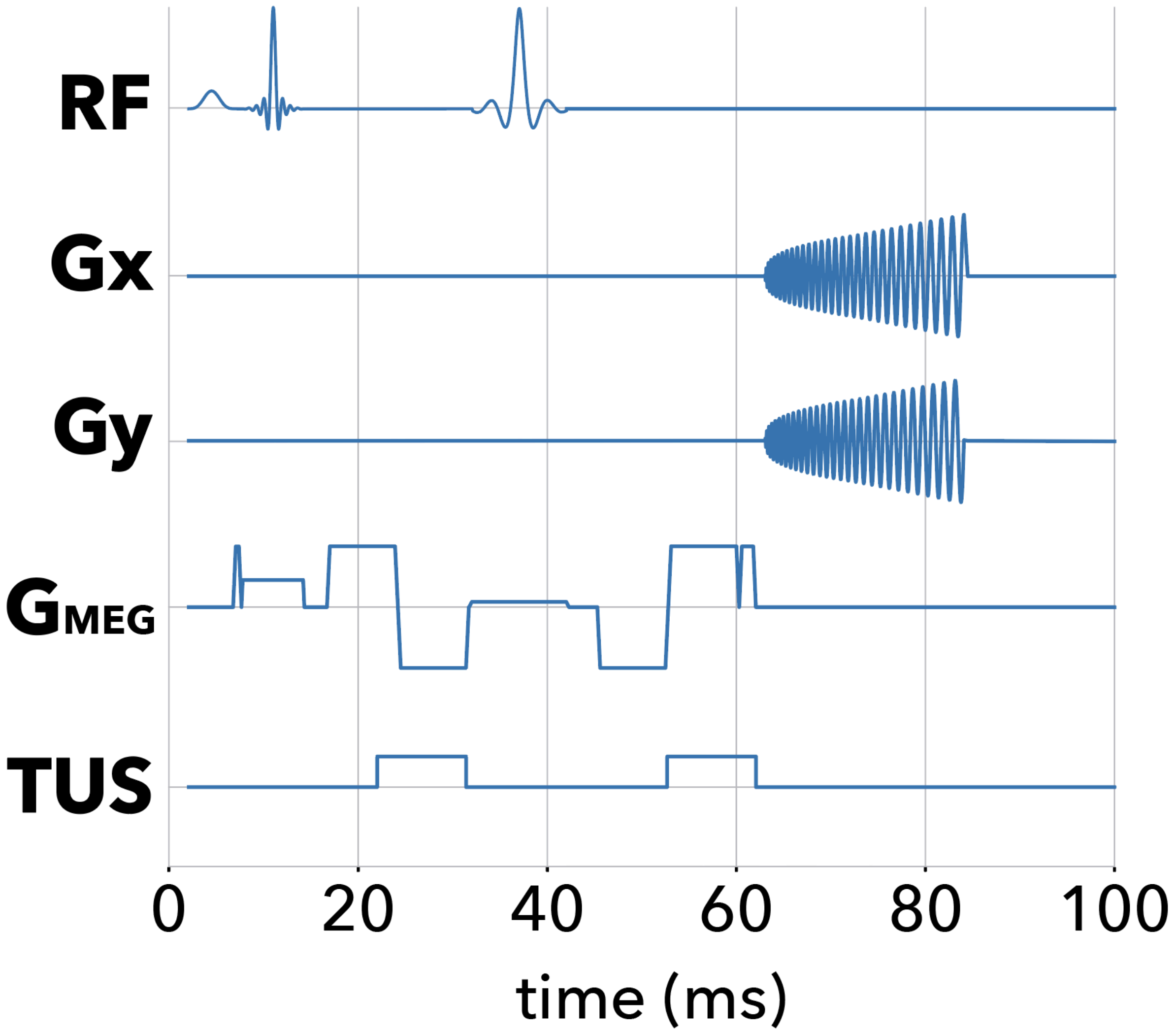


**Supplementary Fig. 8: MR-ARFI pulse sequence.** Repeated inverted bipolar motion encoding gradients were used for phase encoding tissue displacement induced by two 7 ms ultrasound pulses time-locked to begin 2 ms prior to the 5 ms second lobe of each bipolar MEG. This sequence is a motion-robust spin-echo single-shot spiral time series.

#### Supplementary Figure 9 - Counterbalancing

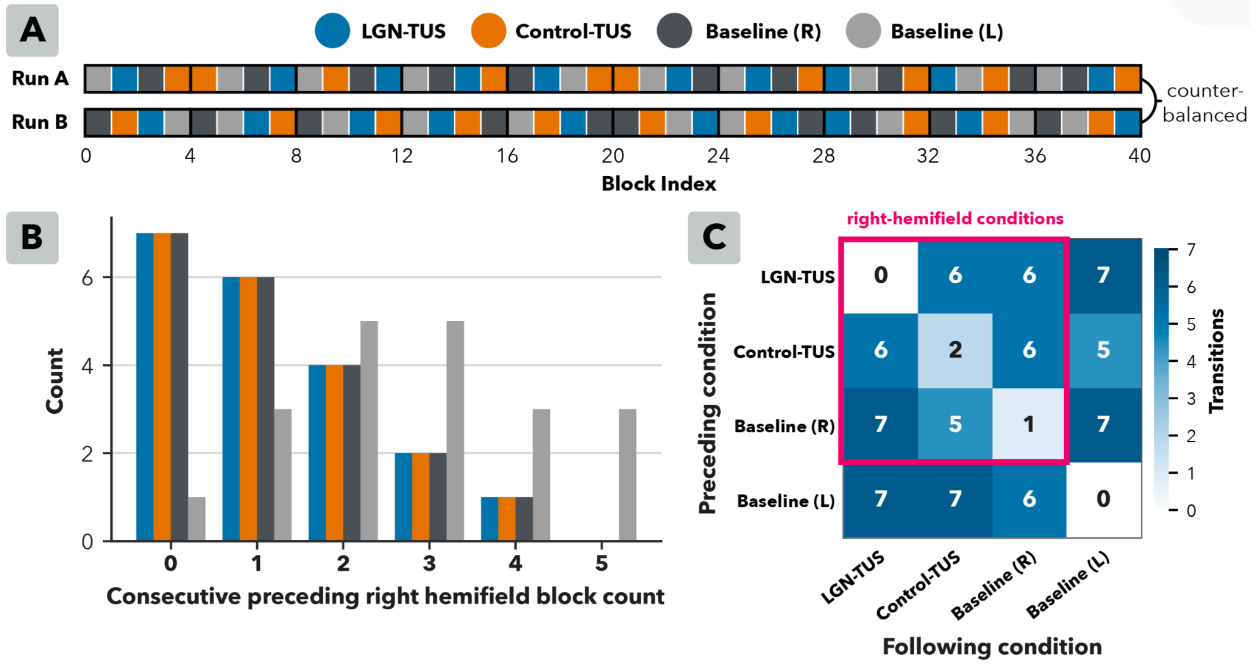


**Supplementary Fig. 9: Condition counterbalancing. (A)** Participants completed two ten-minute functional scans, each comprised of forty fifteen-second blocks. Each 15-second block tested one of four conditions: TUS applied to the LGN (LGN-TUS), TUS applied to the temporal grey matter active control site (Control-TUS), a no-TUS baseline condition with a right hemifield checkerboard stimulus, and a no-TUS baseline condition with a left hemifield checkerboard as a quality control. Each 15-second condition occurred once every four blocks (i.e., dispersed equally across the scan time). Two fixed pseudorandomized orders were generated for the two scans (i.e., Run A and Run B), and the order in which these were acquired was counterbalanced across participants. No more than two ultrasound blocks occurred consecutively to manage heating. Within each ultrasound block, six 145 ms sonications were delivered, time-locked to threshold-level contrast decrement presentation **(B)** For each condition, we equalised visual-adaptation history such that the length of the preceding consecutive right-hemifield visual stimulus blocks had the same distribution across the primary right-hemifield conditions. Condition differences can therefore not be attributed to unequal prior visual adaptation. **(C)** Among the primary right-hemifield conditions, the TUS conditions precede each other condition equally often to balance any carryover effects, and the remaining conditions were equalised in their transitional structure insofar as possible given the ordering constraints specified above.

#### Supplementary Figure 10 - Baseline blinding

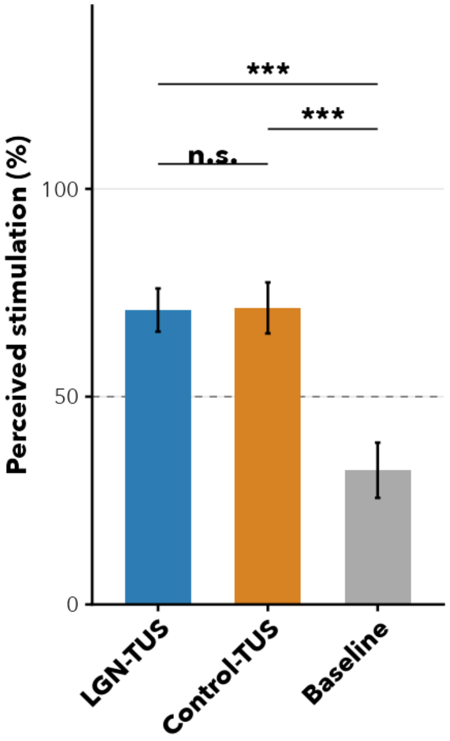


**Supplementary Fig. 10: Blinding of baseline condition.** Although blinding was successful for the primary LGN- vs. Control-TUS comparison, participants were able to distinguish both TUS conditions from baseline, with perceived stimulation rated higher for each (LGN-TUS: $t(23) = 4.53$, $p < 0.001$, MΔ = 38.5% [95% CI: 20.9, 56.1], $d_z = 0.92$; Control-TUS: $t(23) = 4.90$, $p < 0.001$, MΔ = 39.1% [95% CI: 22.6, 55.6], $d_z = 1.00$). This most likely reflects the auditory white noise and 4 kHz tone being insufficient to mask the auditory confound present in both TUS conditions, consistent with the pneumatic earbuds' inability to reproduce the complex harmonics needed to replicate that confound for this protocol and transducer. Alternatively, participants may have detected somatosensory co-stimulation[18] that distinguished TUS from no-TUS trials, though they did not consistently report peripheral somatosensory sensations when asked explicitly during debriefing.

#### Supplementary Fig. 11 - Acoustic simulations

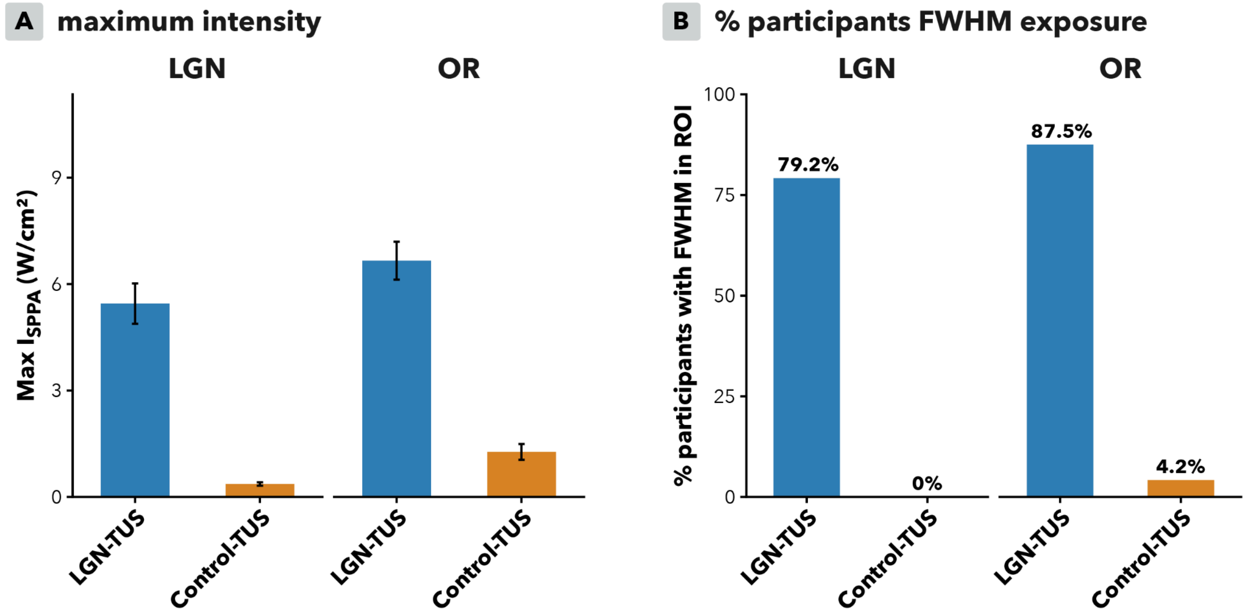


**Supplementary Fig. 11: Acoustic simulations. (A)** The simulated maximum spatial-peak pulse-average intensity ($I_{SPPA}$) in the LGN was 5.45 ± 0.57 W/cm$^2$ (mean ± SE) during LGN-TUS, and only 0.37 ± 0.05 W/cm$^2$ during Control-TUS, confirming condition-specific LGN exposure. Notably, the optic radiation (OR) was also exposed during LGN-TUS (6.66 ± 0.53 W/cm$^2$; Control-TUS: 1.27 ± 0.22 W/cm$^2$). N = 24. **(B)** The full-width-half-maximum (FWHM) of the in-situ simulated focus targeted the LGN in 79.2% of participants during LGN-TUS, and in no participants during Control-TUS. The optic radiation fell within the simulated FWHM in 87.5% of participants during LGN-TUS, and in a single participant in Control-TUS (4.2%). N = 24.

### References


1. Kastner, S. *et al.* Functional Imaging of the Human Lateral Geniculate Nucleus and Pulvinar. *J. Neurophysiol.* **91**, 438–448 (2004).
2. Denison, R. N., Vu, A. T., Yacoub, E., Feinberg, D. A. & Silver, M. A. Functional mapping of the magnocellular and parvocellular subdivisions of human LGN. *NeuroImage* **102**, 358–369 (2014).
3. Schwartz, E. L. Computational anatomy and functional architecture of striate cortex: A spatial mapping approach to perceptual coding. *Vision Res.* **20**, 645–669 (1980).
4. Alvarez, I., De Haas, B. A., Clark, C. A., Rees, G. & Schwarzkopf, D. S. Comparing different stimulus configurations for population receptive field mapping in human fMRI. *Front. Hum. Neurosci.* **9**, (2015).
5. Bayram, A., Karahan, E., Bilgiç, B., Ademoglu, A. & Demiralp, T. Achromatic temporal-frequency responses of human lateral geniculate nucleus and primary visual cortex. *Vision Res.* **127**, 177–185 (2016).
6. Mullen, K. T., Thompson, B. & Hess, R. F. Responses of the human visual cortex and LGN to achromatic and chromatic temporal modulations: An fMRI study. *J. Vis.* **10**, 13 (2010).
7. Schneider, K. A., Richter, M. C. & Kastner, S. Retinotopic Organization and Functional Subdivisions of the Human Lateral Geniculate Nucleus: A High-Resolution Functional Magnetic Resonance Imaging Study. *J. Neurosci.* **24**, 8975–8985 (2004).
8. Schneider, K. A. & Kastner, S. Effects of Sustained Spatial Attention in the Human Lateral Geniculate Nucleus and Superior Colliculus. *J. Neurosci.* **29**, 1784–1795 (2009).
9. Jenkinson, M., Beckmann, C. F., Behrens, T. E. J., Woolrich, M. W. & Smith, S. M. FSL. *NeuroImage* **62**, 782–790 (2012).

10. Aubry, J.-F. *et al.* ITRUSST consensus on biophysical safety for transcranial ultrasound stimulation. *Brain Stimulat.* **18**, 1896–1905 (2025).

11. Treeby, B. E. & Cox, B. T. k-Wave: MATLAB toolbox for the simulation and reconstruction of photoacoustic wave fields. *J. Biomed. Opt.* **15**, 021314 (2010).

12. Miscouridou, M., Pineda-Pardo, J. A., Stagg, C. J., Treeby, B. E. & Stanziola, A. Classical and Learned MR to Pseudo-CT Mappings for Accurate Transcranial Ultrasound Simulation. *IEEE Trans. Ultrason. Ferroelectr. Freq. Control* **69**, 2896–2905 (2022).

13. Zadeh, A. K. *et al.* Enhancing transcranial ultrasound stimulation planning with MRI-derived skull masks: a comparative analysis with CT-based processing. *J. Neural Eng.* **22**, 016020 (2025).

14. Matilainen, N., Kataja, J. & Laakso, I. Verification of neuronavigated TMS accuracy using structured-light 3D scans. *Phys. Med. Biol.* **69**, 085004 (2024).

15. Chen, M. *et al.* Numerical and experimental evaluation of low-intensity transcranial focused ultrasound wave propagation using human skulls for brain neuromodulation. *Med. Phys.* **50**, 38–49 (2023).

16. Farboud, S. *et al.* Rapid modulation of choice behavior by ultrasound on the human frontal eye fields. *Nat. Commun.* **17**, 2966 (2026).

17. Meijer, S. *et al.* The human amygdala in threat learning and extinction. *Sci. Adv.* https://doi.org/10.1126/sciadv.aea8233 (2026) doi:10.1126/sciadv.aea8233.

18. Kop, B. R., de Jong, L., Kim, B. P., den Ouden, H. E. M. & Verhagen, L. Parameter optimisation for mitigating somatosensory confounds during transcranial ultrasonic stimulation. *Brain Stimulat.* **18**, 1224–1236 (2025).
